# Local synaptic inhibition mediates cerebellar granule cell pattern separation necessary for learned sensorimotor associations

**DOI:** 10.1101/2022.05.20.492839

**Authors:** Elizabeth A. Fleming, Greg D. Field, Michael R. Tadross, Court Hull

## Abstract

The cerebellar cortex plays a key role in generating predictive sensorimotor associations. To do so, the granule cell layer is thought to establish unique sensorimotor representations for learning. However, how this is achieved and how granule cell population responses contribute to behavior have remained unclear. To address these questions, we have used *in vivo* calcium imaging and granule cell-specific pharmacological manipulation of synaptic inhibition in awake, behaving mice. We find that inhibition sparsens and thresholds sensory responses, limiting overlap between sensory ensembles and preventing spiking in many granule cells that receive excitatory input. Moreover, we find that inhibition can be recruited in a stimulus-specific manner to powerfully decorrelate multisensory ensembles. Consistent with these results, we find that granule cell inhibition is required for sensory discrimination in a cerebellum-dependent behavior. These data thus reveal new mechanisms for granule cell layer pattern separation beyond those envisioned by classical models.

## Introduction

Associative learning is an essential process linking sensation and action, providing a key mechanism to modify behavior. The cerebellum plays a central role in associative sensorimotor learning, both for generating coordinated movements and cognitive processes^1, 2^. To do so, the cerebellum receives excitatory mossy fiber input from diverse sources^3^ that transmit sensory, motor, and cognitive information to granule cells in the granule cell layer^4^. Granule cells must integrate and relay these signals to Purkinje cells, the output neurons of the cerebellar cortex, in a manner that establishes unique sensorimotor representations necessary for associative learning and the expression of learned cerebellum-dependent behaviors.

Classical models speculate that unique granule cell representations are generated through a process of “pattern separation”^5, 6^. Expansion recoding is one mechanism thought to enable pattern separation, because mossy fiber inputs are distributed onto a population of granule cells that is ∼100-fold larger than the number of mossy fiber inputs^6^. Because each granule cell receives ∼4 inputs that can transmit the same or different modalities^7-10^, random mixing is also thought to facilitate pattern separation. Another key mechanism proposed by classical theories is the thresholding of granule cell activity by local inhibitory interneurons in the granule cell layer called Golgi cells. Golgi cells exhibit spontaneous pacemaker activity, releasing GABA that acts continuously on granule cells to produce a tonic inhibitory current^11, 12^. This tonic inhibition regulates the spike threshold^13^ and gain of granule cell responses^14^. In addition, Golgi cells receive feedforward excitation from mossy fibers and feedback excitation from granule cells, thus allowing them to respond dynamically to the inputs and outputs of the granule cell layer. Together, this tonic and phasic Golgi cell inhibition has long been hypothesized as necessary for creating sparse, non-overlapping granule cell population codes.

In contrast with predictions of classical theories, modern calcium imaging approaches have shown that granule cell responses can be dense and redundant in some conditions^15-18^. These studies have indicated that complex behaviors requiring task engagement, learning, and compound body movements likely to involve sensory, motor, cognitive, and efference copy signals can result in relatively widespread granule cell activity. In such cases, where there are many complex granule cell representations evolving across time, it has been challenging to disentangle discrete sensory representations and how they combine to form complex multisensory codes that remain dissociable for learning and behavior. Moreover, it has been difficult to test what mechanisms shape these sensory representations, as there has been a lack of tools for acute, cell type-specific manipulations of granule cell GABAergic inhibition. Thus, how the granule cell layer encodes discrete sensory input, how local synaptic inhibition contributes to such representations, and what role granule cell inhibition plays in cerebellum-dependent behavior have remained unclear.

To address these long-standing questions, we have used a new approach that allows *in vivo* measurement of cerebellar granule cell population responses while acutely blocking synaptic inhibition in a cell –type-specific manner. Specifically, we have used multiphoton population imaging and behavior in combination with the DART system^19^ (Drugs Acutely Restricted by Tethering) to acutely block synaptic inhibition onto granule cells. In response to discrete sensory input, we find that granule cell population activity is sparse and can be highly variable in terms of response probability, neural ensemble identity, and response timing across trials. In contrast, acutely blocking synaptic inhibition dramatically enhances stimulus-evoked responses, revealing a large population of previously inactive cells, which suggests thresholding is a key mechanism for sparsifying granule cell population ensembles. In addition, this thresholding establishes separate granule cell populations that can only respond to combined multisensory inputs, a property that would not be possible if ensemble sparsity were determined by inputs alone. Surprisingly, we also find that inhibition can be recruited in a stimulus-specific manner, further enhancing pattern separation by removing cells from multisensory ensembles that are part of unisensory ensembles. In support of our finding that synaptic inhibition plays a central role in granule cell layer pattern separation, we find that blocking inhibition onto granule cells impairs the expression of learned, cerebellum-dependent sensorimotor discrimination. Together, these data reveal multiple distinct computations mediated by GABAergic inhibition onto granule cells that support sensory encoding, pattern separation, and behavior in ways not predicted by classical models.

## Results

### Granule cell synaptic inhibition sparsens population-level sensory representations

To measure sensory-evoked activity in populations of cerebellar granule cells, we performed two-photon calcium imaging of GCaMP6f, which has been shown to report spiking in granule cells *in vitro*^16^. By crossing Ai148^20^ and BACα6Cre-A transgenic mice^21^, we observed dense labeling of granule cells, with only rare off-target labeling of Purkinje cells, which were excluded from imaging analysis (Methods, Supp. Fig. 1). We imaged activity in Crus I of the lateral cerebellum (Fig. 1) in response to auditory stimuli (Fig. 1c-g, pure tones: 1, 5 and 10 kHz at 68 and 72 dB; Supp. Fig. 2) and somatosensory stimuli (gentle orofacial air puffs: 10, 15 and 20 PSI; Supp. Fig. 3). In this region, *in vivo* recordings have identified that granule cells respond to these modalities^9, 22, 23^. Moreover, anatomical tracing has revealed that Crus I is a major cerebellar destination of both auditory and somatosensory inputs^24^.

**Figure 1.**
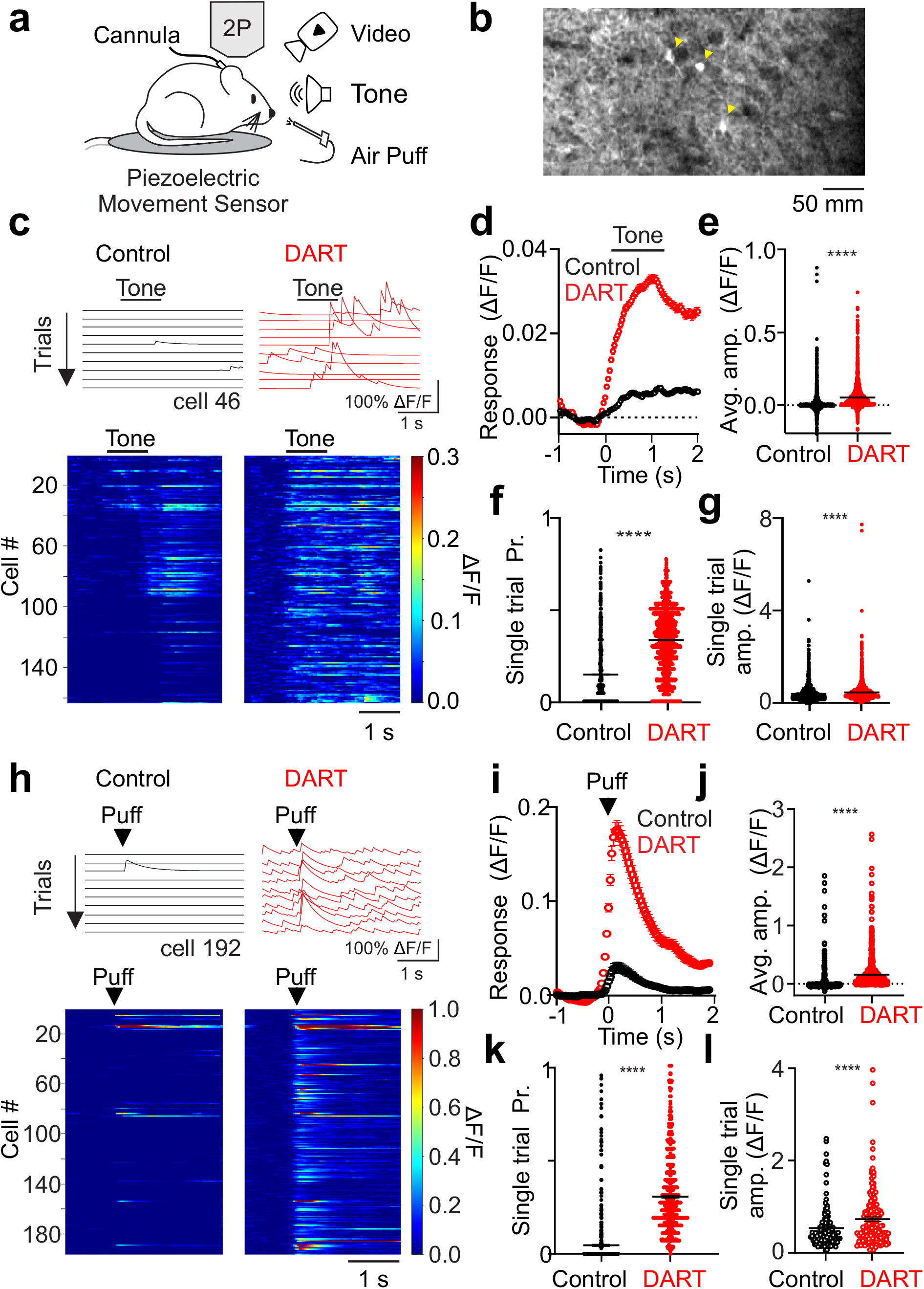
Local synaptic inhibition sparsens and thresholds cerebellar granule cell sensory responses. **a**. Schematic of experimental approach. **b**. Example average field of view across trials of granule cells expressing GCaMP6f during presentation of a somatosensory stimulus in the absence of whole-body movement. Yellow arrows designate significantly responsive cells. **c**. Top, example calcium traces (ΔF/F) from a granule cell on sequential tone presentation trials before (black) and after (red) gabazine.1^DART.2^ infusion (“DART”, 1 μM). Bottom, mean responses of all cells with significant responses to a tone before (left) and after DART application (right) in an example mouse with granule cell HTP expression. Example cell above is cell 46. **d**. Mean time course of responses during tone presentation before (black) and after (red) DART infusion. Error is SEM across cells (n=3360 cells). **e**. Mean response amplitudes for individual cells before (black) and after (red) DART infusion. Black lines are mean ± SEM across cells. **f**. Same as **e**, for response probability. **g**. Same as **e**, for mean responses on all trials with significant responses (n=1942). **h-l**. Same as **c-g**, for responses to somatosensory stimuli (i-k, n=815 cells, l, n=118 cells). Example cell in **h** is cell 192 in the heatmap. \*\*\*\**p* < 0.0001.

Because our goal was to measure discrete sensory responses, and Crus I also receives input related to whisking and likely other movements^25-29^, we took multiple steps to isolate sensory-related granule cell activity. First, animals were habituated to orofacial airpuffs, such that they produced reflexive whisker movements on only a minority of trials. Second, high-speed video was used to detect whisker and facial movements between trials and isolate responses related to these movements^30^ (Supp. Fig. 4). Third, we controlled for spontaneous and reactive body movements using a sensitive piezo vibration sensor^31, 32^ (Figure 1a, Supp. Fig. 5). Together, these methods enabled us to discard trials with body movement and demonstrate that whisking-related activity does not contaminate sensory responses (Supp. Figure 4, 5; Methods).

In control conditions, combining data across stimulus frequencies and intensities reveals that both auditory and somatosensory stimuli recruit population level granule cell activity in Crus I (Fig. 1). All stimuli evoke granule cell responses that begin near the time of stimulus onset (Fig 1c,h). For auditory stimuli, which could be delivered for a longer duration (1 s) than somatosensory stimuli, the mean responses of individual granule cells tile the duration of the stimulus window (Fig. 1c, 4a), and many granule cells respond preferentially at the offset of the stimulus (Fig. 1c). Overall, granule cells in control conditions produce modest responses (Fig. 1d,e,I,j) occurring with a low probability (Figure 1f,k; auditory: 0.14 ± 0.00, n=3360; somatosensory: 0.06 ± 0.00, n=815) and small single trial amplitude (Fig. 1 g, l; auditory: 0.41 ± 0.01 ΔF/F, n = 1942; somatosensory: 0.53 ± 0.04, n = 118). These properties are consistent for all individual auditory and somatosensory stimuli tested (Supp. Fig. 2,3).

Both classical models^5, 6^ and *in vivo* whole-cell recordings^13^ have suggested that inhibition restricts granule cell sensory responses due to spike thresholding. Therefore, local inhibition could explain the low probability and amplitude of individual granule cell trial-by-trial responses. To test how local synaptic inhibition regulates sensory-evoked granule cell responses, we used the DART system^19^ to selectively block GABA_A_ receptors on granule cells (Fig. 1., Supp. Fig. 1,6-8). Here, we expressed a GPI-anchored HaloTag Protein (HTP_GPI_) in granule cells to acutely and specifically antagonize GABA_A_ receptors upon infusion of gabazine.1^DART.2^ (Fig. 1, “DART”). This manipulation dramatically altered responses to all sensory stimuli tested (Fig 1, Supp. Figs. 2,3), producing significantly larger mean population responses (Fig. 1 d-e,i-j), increased response probability (Figure 1f,k; auditory: 0.38 ± 0.00, n = 2220, *P* < 0.0001, paired t-test; somatosensory: 0.31 ± 0.01, n = 727, *P* < 0.0001, paired t-test), and increased amplitude of single trial responses (Figure 1g,l; auditory: 0.60 ± 0.02 ΔF/F, n = 1144, *P* < 0.0001, paired t-test; somatosensory: 0.88 ± 0.07, n = 165, *P* < 0.0001, paired t-test). These changes in single trial response probability and amplitude were observed across all individual variations of auditory and somatosensory stimuli tested (Supp. Fig 2-3).

In addition to changes in sensory evoked responses, we also observed a significant enhancement of spontaneous activity when inhibition was blocked (Fig. 1c,h, control F: 0.000 ± 0.000, DART F: 0.004 ± 0.001, n = 1616, *P* < 0.0001). This effect is consistent with previous data revealing that a non-specific block of GABAergic inhibition in the cerebellar cortex increases spontaneous granule cell spiking and can degrade the signal-to-noise ratio of sensory evoked responses^33^.

To test the selectively of these effects, we used a variation of gabazine.1^DART.2^ that cannot bind HTP (non-binding gabazine.1^nbDART^, “nbDART”) and a variation of HTP that cannot bind ligand (^dd^HTP). Neither nbDART infused into animals expressing HTP, nor DART infused into animals expressing ^dd^HTP significantly changed auditory or somatosensory responses (Supp. Fig. 6,7). Together, these results are consistent with a central role of GABAergic inhibition in enforcing granule spike thresholds to restrict population activity, maintaining sparsity of spiking both within and across trials.

Importantly, many granule cells that are silent in control conditions become responsive after blocking synaptic inhibition, resulting in a dramatic increase in the number of granule cells responsive to both auditory and somatosensory stimulation (Fig. 1c,h, Fig. 2a,b). We therefore compared the number of responsive cells to a conservative estimate of the total cells present in our field of view (based on the size of ROIs detected with our analysis, see Methods). This allowed us to estimate that, under baseline conditions, approximately 5.2 ± 0.5% of granule cells in our field of view responded to any individual auditory stimulus and 1.1 ± 0.2% of granule cells responded to any individual somatosensory stimulus (Figure 2a,b). Following block of synaptic inhibition, there was a large expansion in the fraction of granule cells responsive to both auditory (13.1 ± 1.3%, *P*<0.0001, n = 26, paired t-test) and somatosensory (7.7 ± 2.1%, *P*<0.01, n = 12, paired t-test) stimuli (Fig. 2a,b). Two points merit emphasis: First, these results indicate that the fluorescent granule cell responses we have measured are unlikely to reflect subthreshold activity, as the majority of granule cells were not responsive until inhibition was blocked. These cells necessarily received excitatory input in control conditions that was not reported by GCaMP activity. Second, the sparsity of granule cell population responses we have measured is due to local synaptic inhibition, not a sparsity of incoming mossy fiber input.

**Figure 2.**
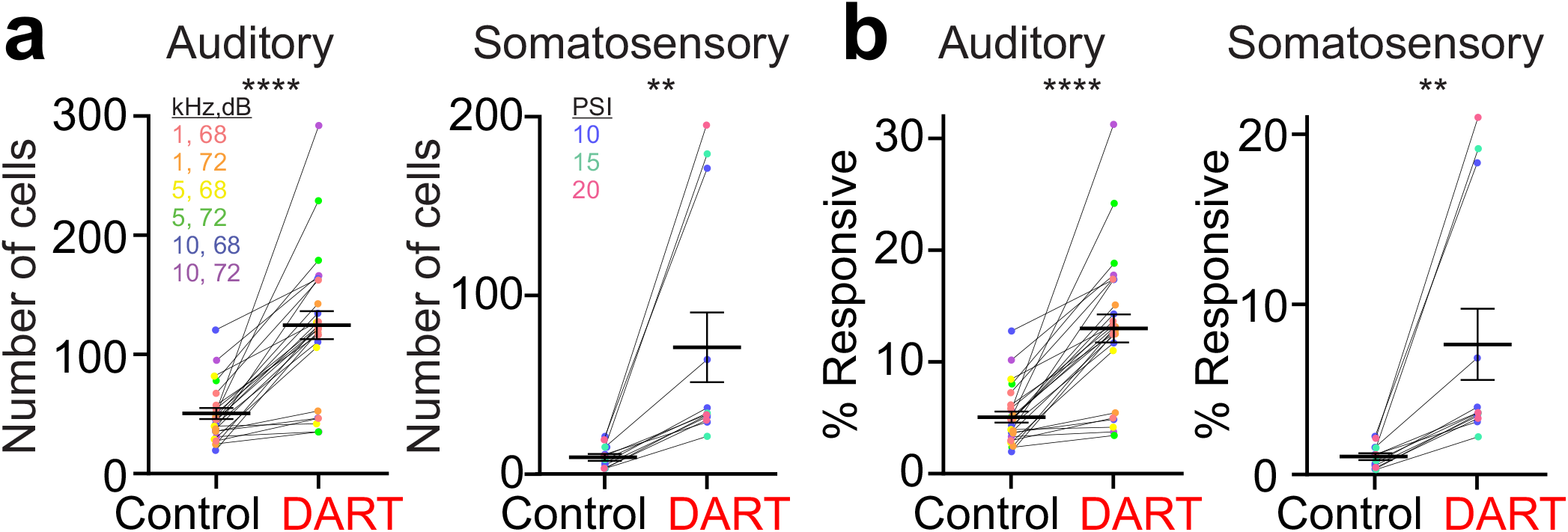
Synaptic inhibition restricts the number of granule cells recruited by sensory input. **a**. Number of responsive cells in each experiment (colors) and across all experiments (black) before and after DART for auditory (left) and somatosensory (right) stimuli. Colors represent specific stimulus conditions; error is SEM across conditions. **b**. Same as **a**, for fraction of total responsive cells.

### Granule cell synaptic inhibition establishes discrete unisensory ensembles via thresholding

Given that each stimulus evokes sparse granule cell activity under control conditions, we next asked whether individual granule cells respond selectively to distinct stimuli. We find that granule cells prefer individual stimulus features, such that cells with robust responses to auditory stimuli of a given frequency respond only weakly to other frequencies (Fig. 3a,c; all comparisons: p<0.0001, one-way ANOVA). Similarly, neurons that respond to a given somatosensory (Fig. 3e; all comparisons: p<0.0001) or auditory (Supp. Fig. 9; all comparisons: p<0.0001) stimulus intensity respond less strongly to other intensities.

**Figure 3.**
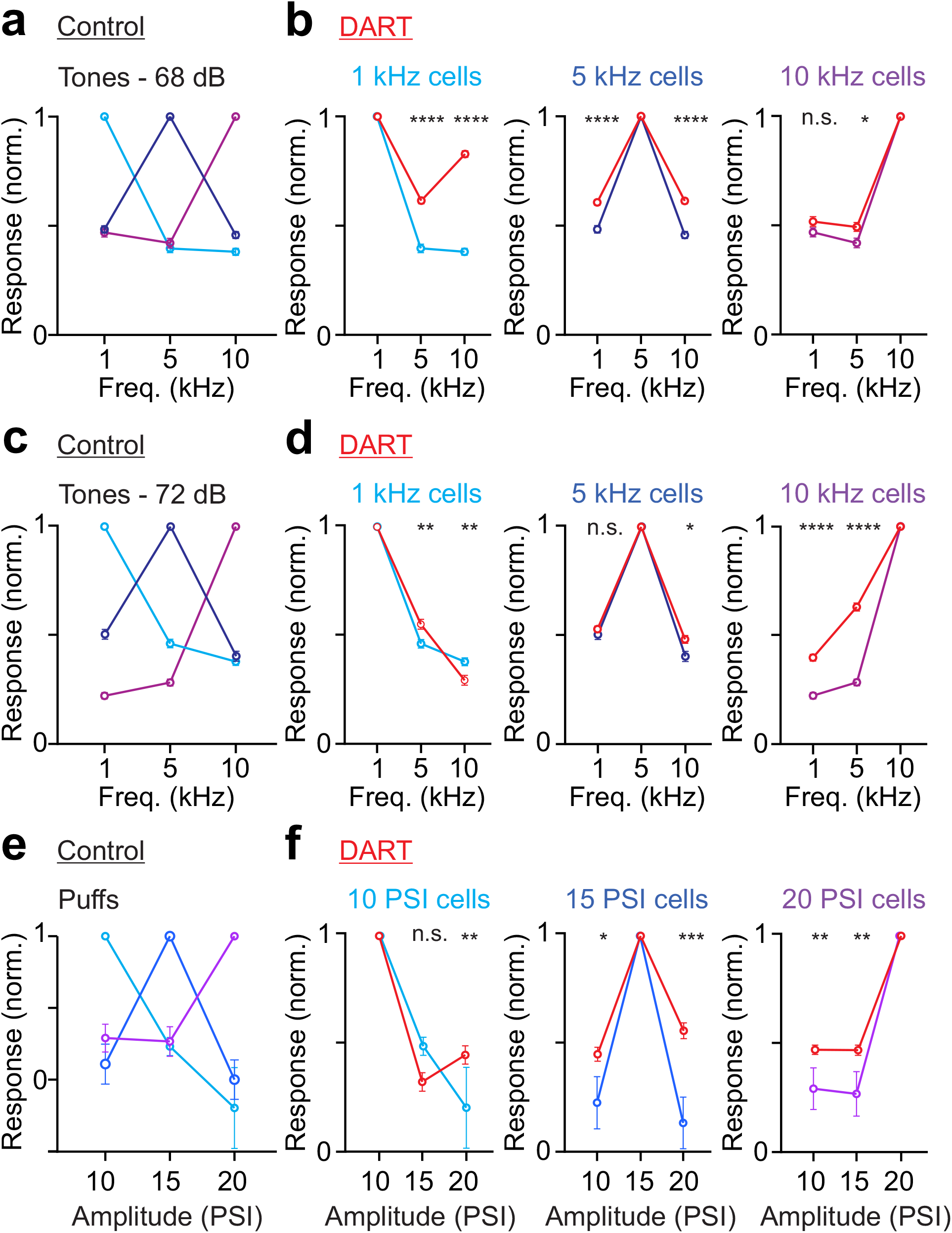
Cerebellar granule cells have stimulus feature preferences that are largely independent of local synaptic inhibition. **a**. Normalized responses in control conditions for all granule cells responsive to 68 dB tones, grouped according to the frequency that drove the maximum response: 1 kHz: n = 227 cells; 3 mice; 5 kHz: n = 211 cells; 10 kHz: n = 166 cells. Error is SEM across cells. **b**. Same as **a**, for tone-responsive granule cells before and after DART infusion (red; 1 kHz: n = 250 cells, 5 kHz: n = 297 cells, 10 kHz: n = 169 cells). **c**. Same as **a**, for granule cells responsive to 72 dB tones preferring: 1 kHz: n = 184 cells; 5 kHz: n = 141 cells; 10 kHz: n = 313 cells. **d**. Same as **b**, for tone-responsive granule cells before and after DART infusion (red; 1 kHz: n = 170 cells, 5 kHz: n = 331 cells, 10 kHz: n = 224 cells). **e**. Same as **a**, for all puff responsive cells preferring: 10 PSI: n = 45 cells; 3 mice; 15 PSI: n = 47 cells; 20 PSI: n = 74 cells. **f**. Same as **b**, for puff responsive cells before and after DART infusion (red; 10 PSI: n = 100 cells, 15 PSI: n = 69 cells, 20 PSI: n = 202 cells).

We next tested how GABAergic inhibition regulates stimulus preferences in granule cells. In neocortical cells, there is evidence that sensory tuning can by sharpened by local synaptic inhibition in some conditions^34^. For granule cells matched between control and after DART infusion (i.e. those responsive in both conditions), we find that removal of synaptic inhibition has a modest effect on stimulus preferences (Fig. 3b,d,f), as mean population preferences remained significant when inhibition was blocked (Supp. Fig. 10; auditory: all comparisons: p<0.0001; somatosensory: all comparisons: p<0.0001). While stimulus tuning was broadened modestly by blocking inhibition, these data suggest that granule cell stimulus preferences are largely inherited from presynaptic mossy fiber input, consistent with the small number of mossy fibers that impinge on individual granule cells and the lack of recurrent processing among granule cells.

To test how these stimulus preferences establish discrete population responses, we first measured the overlap across activated populations. We find that granule cells that respond to each stimulus feature establish ensembles with partial overlap (Fig. 4a,b; auditory fraction overlap: 0.53 ± 0.02, n = 6; somatosensory: 0.06 ± 0.01, n = 3).

**Figure 4.**
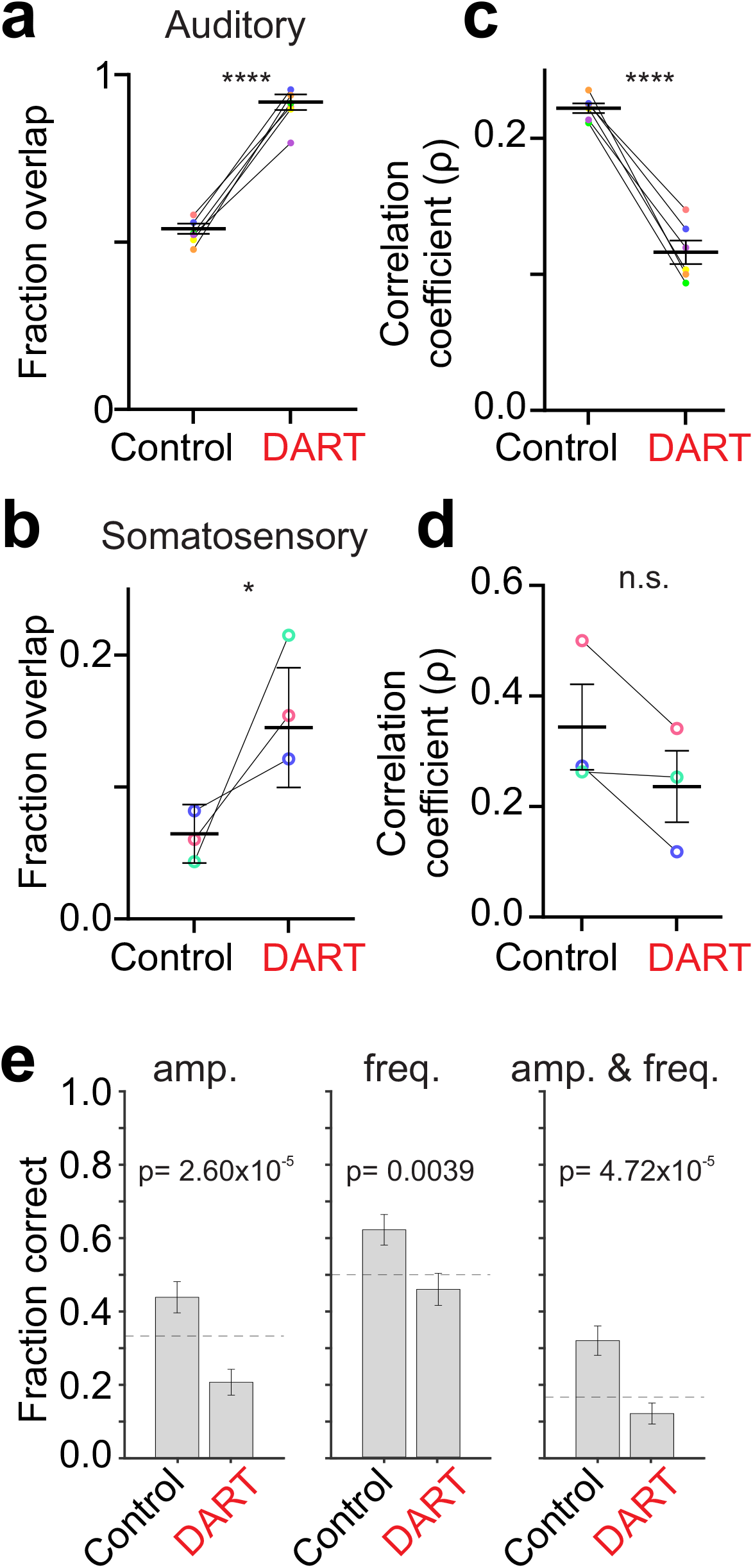
Synaptic inhibition facilitates pattern separation for auditory and somatosensory responsive granule cell ensembles. **a-b**. Fraction of overlap for each stimulus condition (colors) and across all conditions (black) before and after DART for auditory (a) and somatosensory (b) stimuli. Error is SEM across conditions. **c-d**. Same as **a-b** for the Pearson’s correlation of neuronal identity across trials. **c**. Classification performance under control and gabazine.1^DART.2^ conditions for correctly identifying the sound frequency only (left), amplitude only (middle), and amplitude and frequency (right). Dashed lines indicate chance performance. Error bars are S.E.M. estimated using the Wald method for binomial distributions.

We observed less overlap in somatosensory ensembles than auditory ensembles. However, this was likely due to the smaller size of somatosensory ensembles. On average, auditory ensembles contained 83.7 ± 14.7 cells, (n = 18), whereas somatosensory ensembles contained only 29.4 ± 6.5 cells (n = 9). Therefore, the average overlap for auditory ensembles was 44.3 ± 0.2 cells, but only 1.8 ± 0.1 cells for somatosensory ensembles.

In the DART condition, where there is a large number of responsive cells that were unresponsive in control conditions, we find significantly more overlap between the ensembles that are responsive to each individual stimulus (Fig. 4a,b; auditory: 0.90 ± 0.02, n = 6, *P*<0.0001, unpaired t-test; somatosensory: 0.16 ± 0.03, n = 3, *P*=0.03, unpaired t-test). Thus, blocking synaptic inhibition decreases the separability of the average population response to different sensory stimuli.

Sensory discrimination can also be influenced by trial-over-trial variability. Despite representing different stimuli with discrete ensembles, there was significant variability across trials within each ensemble of cells responding to any given stimulus feature (Fig. 4c,d). On average, each ensemble of responsive granule cells was only weakly correlated across trials (auditory: correlation coefficient (ρ = 0.22 ± 0.00; somatosensory: ρ = 0.34 ± 0.08). Thus, while the sparse population response allows for discrete granule cell ensembles to represent individual stimulus features on average, the ensembles are surprisingly stochastic across trials, a feature that may contribute to the relatively slow time course of most cerebellar learning.

Importantly, the large number of newly active cells in DART also added variability to ensemble identities, as ensembles became even less correlated across trials when inhibition was blocked (Fig. 4c,d; auditory: 0.12 ± 0.01, 21.9 ± 0.8 trials/ensemble, *P*<0.0001, unpaired t-test; somatosensory: 0.24 ± 0.06, 16.7 ± 2.9 trials/ensemble, *P*=0.35, unpaired t-test). These results suggest that the higher mean response probability in DART (Fig. 1) is not sufficient to counter the variability introduced by the large number of additional responsive cells when inhibition is blocked. Together, these results indicate that inhibition serves to segregate granule cell ensembles representing discrete stimulus features largely by thresholding population activity.

To test whether the population response can discriminate the different stimuli presented despite the relatively high level of trial-to-trial variability, we trained a decoder to identify the stimulus classes that produced granule cell population responses (Methods). Under control conditions, using a population of 321 granule cells, single trial auditory responses could be correctly categorized above chance according to their amplitude, their frequency, or the combination of amplitude and frequency (Fig. 4e, amplitude p=0.008, frequency p=0.003, amplitude and frequency p=6.26×10^−5^). When synaptic inhibition was blocked with DART, however, categorization was significantly impaired for all stimulus conditions (Fig. 4e-h, control vs DART; amplitude p=2.60×10^−5^, frequency p=0.004, amplitude and frequency p = 7.72×10^−5^), falling below chance performance. For somatosensory responses, the smaller ensemble sizes prevented robust decoding, but the same trends remained comparing control and DART conditions (Control: 51.1 ± 12.9% correct DART: 42.2 ± 12.8% correct (chance = 33% correct), p = 0.2). Together, these results demonstrate a key role for granule cell synaptic inhibition in sensory pattern separation..

### Granule cell synaptic inhibition establishes discrete multisensory ensembles via multiple computations

Our data indicate that excitatory inputs from mossy fibers and local synaptic inhibition can establish distinct ensembles of granule cells that encode individual sensory stimuli. However, it has also been demonstrated anatomically and physiologically that some granule cells receive mossy fiber input from more than one source^7-9^, and it is thought that integration of these inputs can enhance the diversity of granule cell encoding^5-7^. Therefore, we next tested the principles that govern this integration by examining population responses to overlapping stimuli of two different modalities (Figure 5).

**Figure 5.**
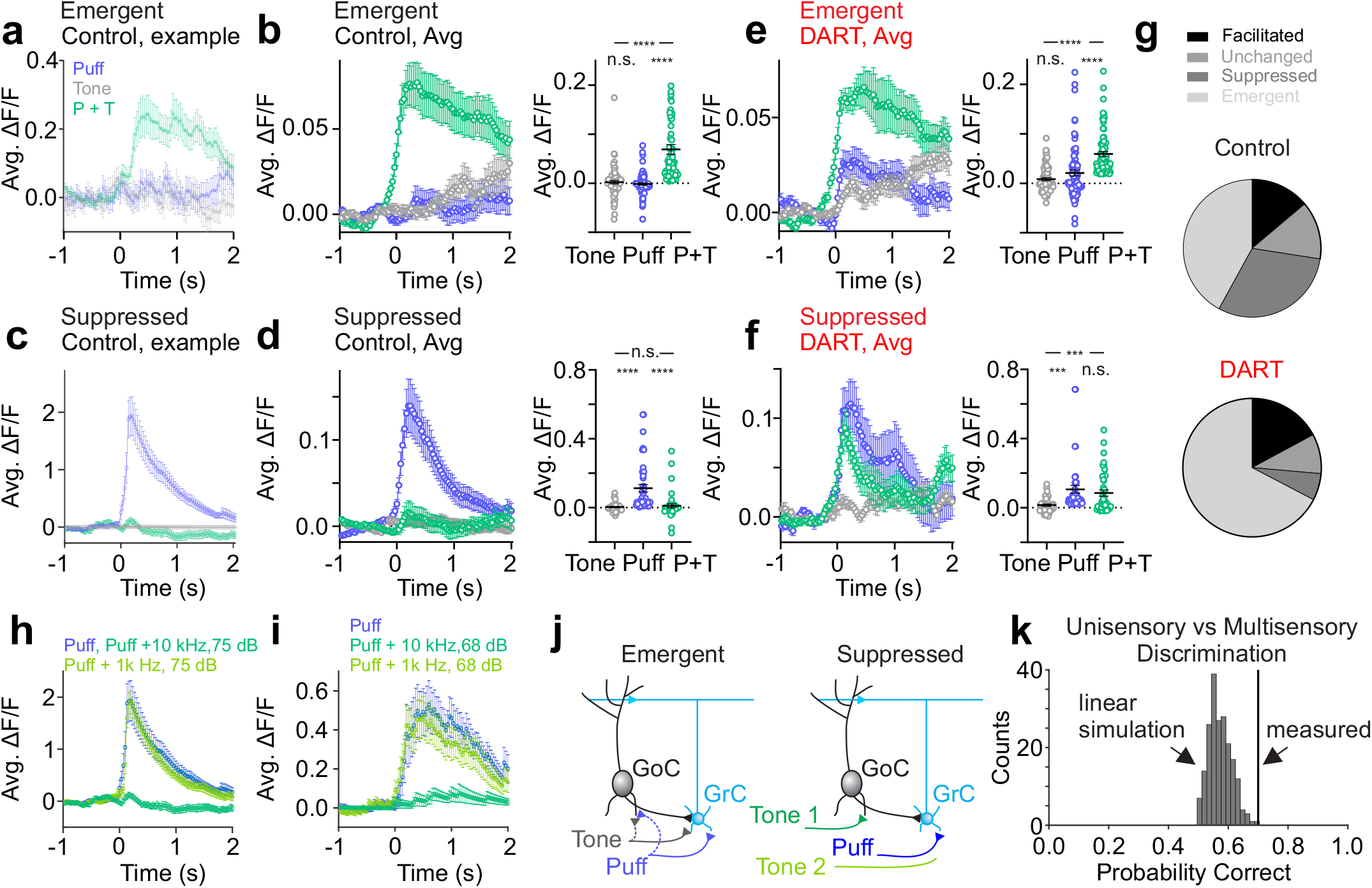
Coincident stimuli create unique granule cell ensembles. **a**. Time course of responses to uni- (tone: gray; puff: blue) and multisensory (tone + puff: green) stimuli for an example emergent cell. Error is SEM across trials. **b**. Left, same as **a**, for all emergent cells (n=79, subset matched to DART condition). Right, amplitude of responses to tone, puff and tone + puff for all matched emergent cells. Error is SEM across cells. **c-d**. Same as **a-b**, for matched suppressed cells (n=43). **e-f**. Same as **b** and **d**, for matched cells after DART infusion. **g**. Pie charts illustrating relative prevalence of each response category before (top; facilitated: n=161, unchanged: n=158, suppressed: n=355, emergent: n=488) and after (bottom; facilitated: n=172, unchanged: n=93, suppressed: n=63, emergent: 676) DART infusion. **h-i**. Example cells illustrating stimulus-specific suppression. Both cells respond robustly to the puff alone (blue), but each is suppressed in response to a specific combination of puff and tone of a given frequency and amplitude (dark green), but not a different frequency and amplitude (light green). Error is SEM across trials. **j**. Granule cell layer circuit motif that would produce emergent responses (left) and stimulus-specific suppression (right). GrC- granule cell; GoC- Golgi cell. \*\*\**p* < 0.001, \*\*\*\**p* < 0.0001. **k**. Black line indicates uni- vs. multisensory discrimination performance from granule cell population responses. Distribution shows discrimination performance for 200 sets of simulated multisensory responses generated by linearly combining random draws from two unimodal responses (one auditory and one somatosensory). The classification performance indicated by the black line has a z-score of 3.21, corresponding to a p-value of 0.0023.

Consistent with previous single cell recordings showing enhanced spiking in response to convergent multisensory mossy fiber input^7, 9^, we find that some cells exhibit larger responses to combined auditory and somatosensory input than to somatosensory input alone (Supp. Fig. 11, “facilitated”) (n = 161, *P* < 0.0001, puff vs. tone + puff: *P* < 0.0001, RM ANOVA with Tukey’s multiple comparisons). Thus, part of the population code representing multisensory stimulus combinations is reflected by increased activity within the same cells that respond to each stimulus in isolation.

We also find, however, that the identity of cells that define multisensory ensembles differ with respect to the ensembles representing each stimulus in isolation in two important ways: First, we find that many granule cells with no significant responses to either stimulus alone became active in response to combined auditory and somatosensory stimulation (Figure 5a,b,g, “emergent”) (n = 488, tone or puff vs. tone + puff: *P* < 0.0001, RM ANOVA with Tukey’s multiple comparisons). These data suggest that integration of both stimulus modalities is necessary to drive these granule cells above spike threshold. These ‘emergent’ cells thus generate a novel multisensory ensemble by adding new cells that are not present in the ensembles representing each stimulus in isolation.

Second, we also observed a large population of granule cells that are suppressed in response to combined somatosensory and auditory stimulation (Figure 5c,d,g, “suppressed”) (, n = 355, puff vs. tone + puff: *P* < 0.0001, RM ANOVA with Tukey’s multiple comparisons). Many of these cells are completely silenced, thus subtracting them from the ensembles representing individual stimuli in isolation. This effect therefore further separates the new, multisensory ensemble from the unisensory ensembles. Together, the emergent and suppressed populations of granule cells represent the vast majority of total granule cells in our measurements (Figure 5g), suggesting that whatever mechanism is responsible for these computations is critical for the encoding of complex multisensory stimuli in the granule cell layer.

To test whether local synaptic inhibition provides the necessary mechanism to establish suppressed and emergent populations, we again utilized the DART system to acutely block inhibition. An analysis of matched cells between control and DART conditions revealed that inhibition powerfully shapes both the emergent and suppressed granule cell populations (Fig. 5e,f,g). Specifically, though emergent cells had no significant response to either stimulus alone in control conditions, we find that blocking synaptic inhibition revealed responses to each individual stimulus (Fig. 5e)(n = 79, tone = 0.8 ± 2.9% ΔF/F, puff = 2.1 ± 0.6% ΔF/F, *P*<0.0001, paired t-test), again consistent with a spike thresholding effect. These data also contextualize the strategy of utilizing widespread subthreshold input instead of sparse, high-fidelity suprathreshold input, as it would not be possible to generate these unique emergent multisensory ensembles with the latter strategy.

We also find that the suppressed population was dependent on synaptic inhibition, as suppression was abolished in the presence of DART (Fig. 5f) (n = 43, puff vs. tone + puff: *P* = 0.38, RM ANOVA with Tukey’s multiple comparisons). These data indicate that inhibition can be recruited in a stimulus-specific manner, where in this case, auditory input can recruit inhibition that suppresses responses to somatosensory input. In this manner, local inhibition can mediate subtractive stimulus integration, operating to suppress the response of one input when another is present. Consistent with theoretical work, which has hypothesized that such a mechanism could act to diversify granule cell representations by reducing ensemble overlap during combined stimulus presentations^35^, our data indicate that stimulus-specific suppression provides a widespread and powerful means to generate unique multisensory ensembles.

To further explore this computation, we varied the features of co-presented stimuli (Fig. 5h,i,j). We find that suppression can be specific even within the same modality during co-presentation of stimuli with different features (Fig. 5h,i). These data suggest that local synaptic inhibition is a crucial source for generating diversity in granule cell population responses during complex sensory input, segregating multisensory ensembles from each other and from those representing each stimulus in isolation.

To test this directly, we again trained a decoder to categorize a stimulus as either unisensory (auditory or somatosensory), or multisensory. Stimuli were correctly categorized on 70% of trials (Fig. 5k, decoding 521 granule cell responses from 30 unisensory and 30 multisensory trials). To determine how stimulus-specific inhibition and thresholding of emergent cells contribute to discrimination of multisensory ensembles, we synthesized multisensory population responses that consisted exclusively of the linear summation of measured unisensory responses (Methods). We find that these synthetic multisensory responses are significantly less discriminable from the unisensory responses as compared to measured multisensory responses (Fig. 5k, p = 0.0023). This reveals that the population diversity generated by the novel multisensory interactions identified here (stimulus-specific suppression and emergent responses) serve to significantly enhance pattern separation in the granule cell layer.

### Granule cell sensory responses are temporally variable

In addition to population identity, the timing of granule cell activity has been hypothesized to play an important role in behavior and learning^4, 36, 37^. Therefore, we also tested how granule cells represent the timing of sensory input and how this depends on local synaptic inhibition. We find that the average responses of individual granule cells during a 1-second auditory stimulus forms a population response that tiles the duration of the stimulus (Fig. 6a)^38^. Surprisingly, however, we find that granule cells do not respond with reproduceable timing across trials (Fig. 6a). Because the peak of mean ΔF/F responses can be biased by a small number of trials with large responses, we also computed response timing according to the onset of fluorescent responses during the stimulus window on individual trials (Fig. 6b,c). This measure again supports the conclusion that the timing of granule cell responses is not reproduceable across trials (Fig. 6c). To quantify this variability, we measured the jitter in response onset across trials (Fig. 6d) and found that granule cells have an onset time that varies by hundreds of milliseconds across trials (Fig. 6f; control onset S.D. = 0.77 ± 0.02). These results suggest that individual granule cells do not exhibit a high degree of across-trial consistency, even for a relatively short 1-second stimulus, as has been thought necessary for cerebellar learning^39^.

**Figure 6.**
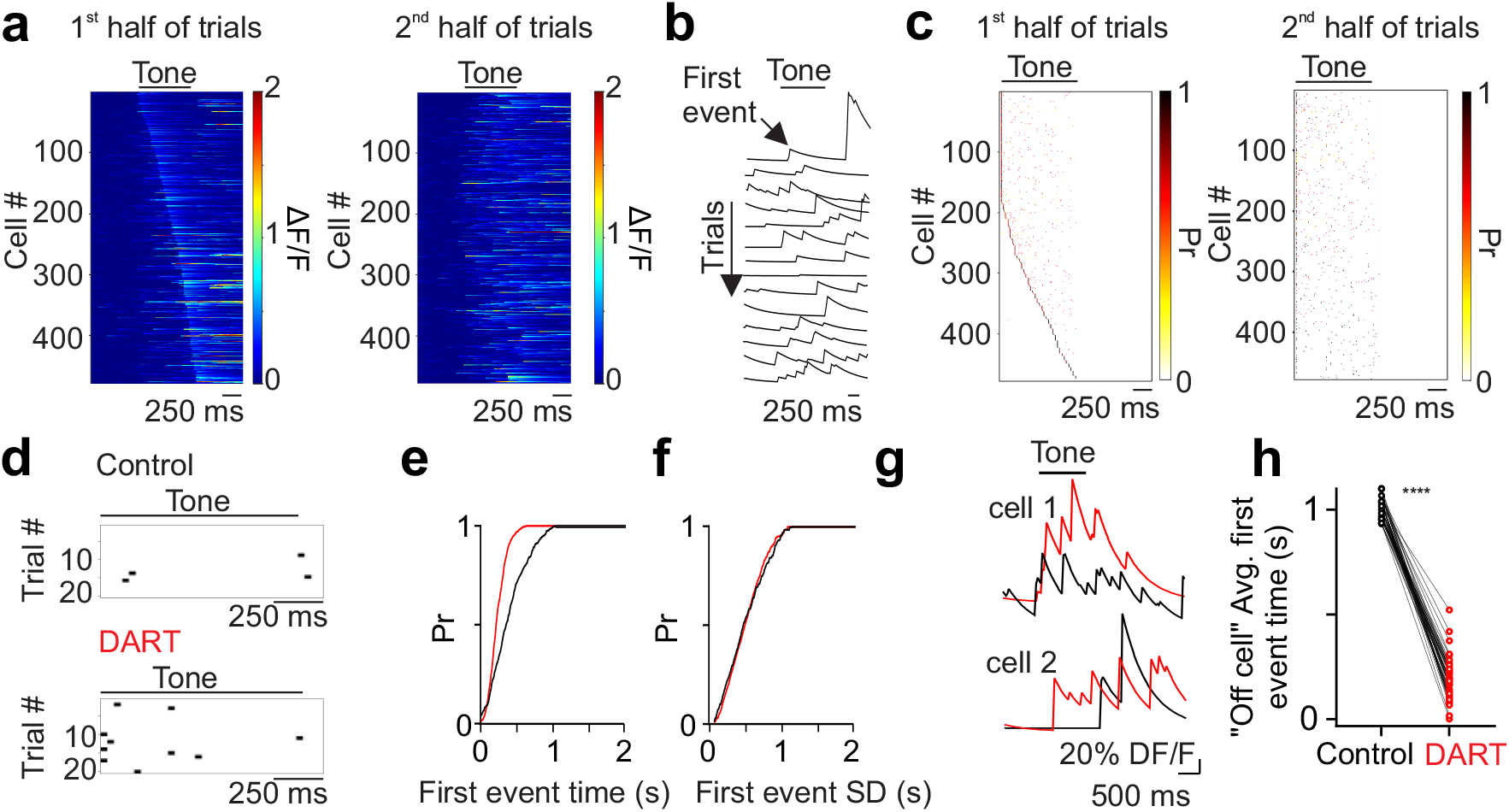
Cerebellar granule cells respond with temporal variability to auditory stimuli. **a**. Left, mean tone-evoked responses on the first half of trials for all cells ordered by peak response time (n = 478 cells). Right, same as on left for the second half of trials, ordered according to peak responses during the first half. **b**. Example calcium traces (ΔF/F) from a granule cell during sequential tone presentation trials. Arrow notes first event following tone onset. **c**. Left, heatmaps displaying the mean first event time probabilities for all cells during the first half of trials ordered by timing of greatest first event probability. Right, first event time probabilities during the second half of trials ordered the same as on left. **d**. Heatmaps displaying peak event times for all significant trials for an example cell in control (top) and after DART infusion (bottom). **e**. Cumulative distribution of mean first event times for all cells in control (black; n=1631) and after DART infusion (red; n=3718; Kolmogorov-Smirnov *P*<0.0001). **f**. Same as **e**, for mean standard deviation of first event times (Kolmogorov-Smirnov *P*=0.9859). **G**. Example traces in control (black) and after DART infusion (red) for two example cells. Note the cell on the top has an increase in peak activity without a change in onset, while the cell on the bottom responds to tone offset in control but tone onset in DART. **H**. Mean response time in control (black) and after DART (red) for cells with a mean first peak time at or after tone offset in control conditions (n=47; paired t-test *P*<0.0001).

We find that blocking synaptic inhibition shifts the distribution of response onset times earlier (Fig. 6e and Supp. Fig. 12-13; first peak: control = 391.2 ± 11.5 ms, DART = 235.6 ± 5.5 ms, n=478, *P*<0.0001, paired t-test). Surprisingly, however, we do not observe a corresponding reduction in response jitter (Figure 6f, DART onset S.D. = 0.77 ± 0.02, *P*=0.59, paired t-test). This was due to the higher response probability across trials in the DART condition, where there were many more total trials with significant stimulus responses (Fig. 6d). As a result, while there were more trials with earlier response times when inhibition was blocked, there were also more trials with late responses, preventing an overall change in mean response jitter (Figure 6d,f). These data are consistent with previous results suggesting that response timing in granule cells is not exclusively regulated by synaptic inhibition^40^ and must, therefore, also reflect parameters such as variability in input timing, and cellular properties such as short-term plasticity of mossy fiber input^7^ and the intrinsic membrane properties of granule cells^41^.

In control conditions, we also find that many granule cells respond at or near the offset of the auditory stimulus (Fig. 1c, 6a,g). These ‘off’ cells represented 5.4% of the responsive population. When inhibition was blocked, these cells responded much earlier during the auditory stimulus (Fig. 6g-h; n = 47, *P*<0.0001, paired t-test). This indicates that, under control conditions, recruitment of inhibition during the sensory stimulus prevents these cells from responding immediately, even though they receive sufficient excitation during the stimulus to drive spiking if inhibition is removed. Together, these data suggest that inhibition serves to diversify the temporal responses of granule cells by both limiting the number of early responses during a stimulus presentation and establishing a population of cells that selectively represents the late component of sensory input.

### Granule cell synaptic inhibition is required for cerebellum-dependent sensory discrimination

Our data reveal that synaptic inhibition powerfully restricts the population of granule cells recruited by sensory input and shapes many features of the population ensembles that encode sensory stimuli. In principle, these effects may support the predictions of classical and recent computational models^5, 6, 42^ proposing that granule cell inhibition acts to segregate the population codes necessary for both learning and the expression of learned behaviors. To test this hypothesis directly, we utilized the cerebellum-dependent task delay eyelid conditioning. For this task, mice were trained on a freely moving treadmill^43^ to associate a brief corneal air puff (unconditioned stimulus, “US”) with a co-terminating neutral auditory tone (5 kHz, 250 ms; conditioned stimulus, “CS+”; Fig. 7a-d). We first trained mice until the probability of conditioned responses (“CR”) in each session had plateaued (0.80 ± 0.06, n = 8; Fig. 7d). In a subset of mice, infusion of muscimol into the eyelid conditioning region of the cerebellar cortex at the floor of the primary fissure, which spread into the anterior interpositus^43^, abolished conditioned responses, confirming cerebellar dependence (n = 3; Fig. 7c and Supp. Fig. 14).

**Figure 7.**
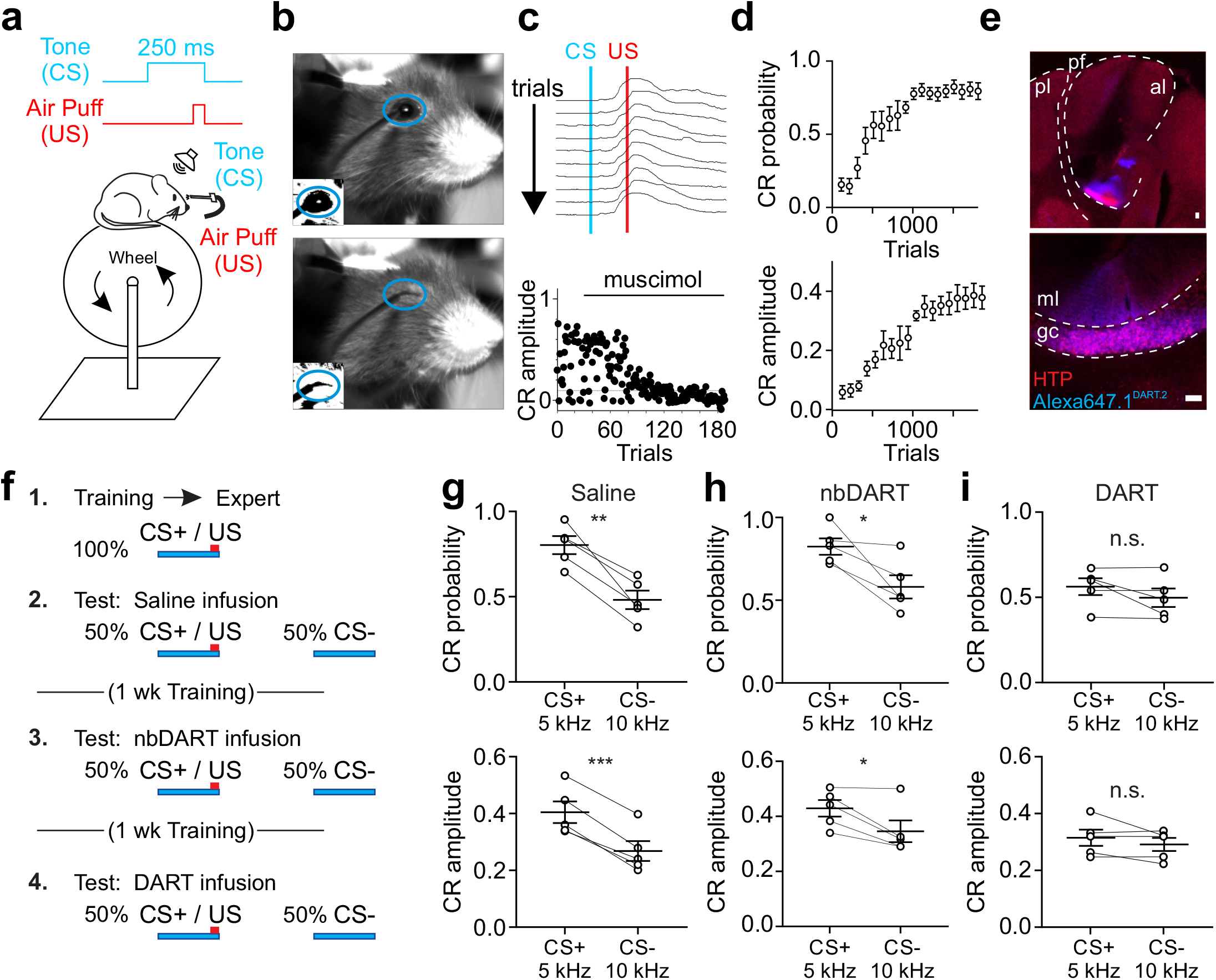
Local synaptic inhibition is necessary for expression of a sensorimotor association. **a**. Schematic of experimental design. CS- conditioned stimulus (tone); US- unconditioned stimulus (corneal air puff). **b**. Sample frames before (top) and after (bottom) air puff delivery. Blue oval indicates ROI analyzed to evaluate conditioned responses. Insets show thresholded binary image used for calculating fractional eyelid closure. **c**. Top, example consecutive eyelid traces from a trained mouse. Blue line mark onset of tone (CS). Red line marks onset of US. Note the eyelid begins to close before the US. Bottom, time course of conditioned response (CR) amplitude for a trained mouse on trials before and after local infusion of muscimol (1 mM). **d**. Average CR probability (top) and amplitude (bottom) during training (n=8 mice). Error is SEM across mice. **e**. Example confocal images (scale 50 μm) of a representative mouse expressing HTP (red) in granule cells in the eyelid conditioning region with bound Alexa647.1^DART.2^ (cyan). Top, pl: posterior lobe; al: anterior lobe; pf: primary fissure. Bottom, ml: molecular layer; gc; granule cell layer. **f**. Experimental time course. CS+: tone frequency paired with air puff (5 kHz); CS-: unpaired tone frequency (10 kHz); nbDART: non-binding gabazine.1^nbDART^. **g**. Average CR probability (top) and amplitude (bottom) for individual mice during presentation of CS+ and CS- trials after saline infusion (n=5 mice). **h**. Same as **g**, after infusion of gabazine.1^nbDART^ (1 μM). **i**. Same as **g**, after infusion of gabazine.1^DART.2^ (1 μM). \**p* < 0.05, \*\**p* < 0.01 \*\*\**p* < 0.001.

To test whether mice could discriminate between the learned CS+ and similar auditory stimuli that had not been paired with a US during learning (CS-), we next measured responses in a cohort of mice for which two different CS-tones (1 and 10 kHz) and the CS+ tone were presented on randomly interleaved trials (Supp. Fig. 15). In these experiments, consistent with previous work in rabbits^44, 45^, we find that the response probability and amplitude of CRs were significantly reduced for both CS-tones as compared to the CS+ tone (Supp. Fig.14; n = 4; CR probability: CS+ vs. CS- (1 kHz or 10 kHz): *P* < 0.005; RM ANOVA with Dunnett’s multiple comparisons; CR amplitude: CS+ vs. CS- (1 kHz or 10 kHz): *P* < 0.05).

To test whether granule cell inhibition is a critical component of sensory discrimination in this cerebellum-dependent task, we again utilized the DART system to block GABA_A_Rs. In a well-trained cohort expressing HTP in granule cells in the eyelid conditioning region of the cerebellar cortex (Fig. 5e), mice were subjected to 3 test sessions, each one week apart with daily CS+ only training in between (Fig. 7f). Test sessions with control infusions of saline or nbDART into the eyelid conditioning region revealed that trained mice effectively distinguished the CS+ from a single tone with a different frequency (10 kHz, 250 ms; “CS-”) (Fig. 7g-h; n = 5; CS+ vs CS- in saline: CR probability: *P* < 0.005, paired t-test; CR amplitude: *P* < 0.0005; CS+ vs CS- in nbDART:, CR probability: CS+ = 0.83 ± 0.05, CS- = 0.58 ± 0.07, n = 5, *P* = < 0.031905, paired t-test, CR amplitude: CS+ = 0.43 ± 0.03, CS- = 0.35 ± 0.04, n = 5, *P* <= 0.031405, paired t-test). However, blocking synaptic inhibition to granule cells in the eyelid conditioning region with functional DART reduced conditioned response probability and amplitude during CS+ trials such that behavior was indistinguishable from CS-trials (Fig. 7i; n = 5; CS+ vs CS- in DART: CR probability: *P* = 0.19, paired t-test; CR amplitude: *P* = 0.25). DART infusion did not significantly impact responses to the CS- (saline vs DART: CR probability: *P* = 0.89, paired t-test; CR amplitude: *P* = 0.69). Interestingly, we did not observe an effect on the trajectory of CRs when granule cell inhibition was blocked (Supp. Fig. 16), suggesting that response kinematics may be largely shaped downstream, perhaps by inhibition and excitation onto Purkinje cells or elsewhere in the circuit. Together, these data indicate that granule cell inhibition is necessary for sensory discrimination in a cerebellum-dependent task and shapes the contextual representations that are harnessed for cerebellar behaviors.

## Discussion

By imaging granule cell responses to discrete sensory input while manipulating local synaptic inhibition, we have revealed multiple key computations that extend classical Marr-Albus models of granule cell layer processing. First, we find that local synaptic inhibition can enforce sparse granule cell population activity in terms of both response probability and cell number. Moreover, consistent with its role as a pattern separation layer, we find that granule cells represent sensory stimuli as discrete ensembles that also depend on inhibition to minimize overlap. For multisensory ensembles, inhibition plays a central role in defining population codes by establishing multisensory-only cells and by suppressing responses of unisensory cells in a stimulus-specific manner. In these ways, which were not envisioned by classical models, inhibition can generate novel multisensory granule cell ensembles that enhance pattern separation. Finally, in agreement with our imaging data suggesting that inhibition serves a central role in establishing discrete sensory representations, we find that granule cell inhibition is required for sensory discrimination in a cerebellum-dependent behavior.

### Granule cell population coding and pattern separation

Recent work has shown that, during complex behaviors, granule cell activity can be denser than was predicted by classical Marr-Albus models^15-18^. In contrast with these experiments, our goal was to isolate discrete sensory responses, independent of motor and other information. With this experimental design, we observed sparse population responses that were dependent on intact synaptic inhibition. In our unisensory stimulus experiments, inhibition acted primarily via spike thresholding, as blocking inhibition dramatically increased the responsive population, producing large scale overlap across ensembles.

Spike thresholding also played a key role in generating multisensory ensembles by establishing cells that only respond to specific input combinations. These multisensory-only cells rely on intact inhibition, as subthreshold inputs from individual stimuli become suprathreshold when inhibition is blocked. In this way, inhibition creates multisensory ensembles that are unique to specific stimulus combinations. Notably, this would not be possible with sparse mossy fiber input that instead drove high fidelity granule cell spiking. Thus, we speculate that it may be more appropriate to consider thresholding inhibition as a mechanism to enhance combinatorial diversity of granule cell responses than to enforce population sparsity per se.

Beyond this broad thresholding effect, however, we also find that inhibition can be recruited in a stimulus-specific manner to provide subtractive refinement of granule cell population activity. This mechanism regulated a large fraction of cells in our multisensory experiments, representing a powerful additional means of pattern separation that has been previously uncharacterized in the cerebellum. Such stimulus-specific lateral inhibition likely provides an explanation for early measurements suggesting that granule layer inhibition can vary according to the specific combination of vestibular afferent nerve stimulation^46^, and is supported by *in vitro* measurements suggesting that Golgi cells can be recruited in a pathway specific manner^47^. Moreover, recent imaging experiments have revealed that, while many Golgi cells are synchronously active during behavior, a large fraction can also display diverse activity patterns that may reflect stimulus specific drive^48^.

Here, our measurements reveal that stimulus-specific recruitment of inhibition is a widespread mechanism that shapes multisensory population activity *in vivo*. Notably, this mechanism is in direct opposition to the concept of Golgi cell inhibition of granule cells acting exclusively as a broad, general feedback system proposed by classical Marr-Albus theories. Thus, for multisensory inputs, the combination of adding new cells via thresholded summation and subtracting cells via stimulus-specific inhibition provides a powerful means for pattern separation.

Together, these findings suggest that inhibition can provide the granule cell layer with an even greater capacity for pattern separation than was envisioned by classical Marr-Albus models, and it may serve to counteract the intrinsic limitations on combinatorial diversity that are imposed by the anatomical architecture of mossy fiber input to granule cells^49^. Moreover, these measurements support the idea that the granule cell layer can generate codes that are high dimensional^50^, in particular, by decorrelating sensory representations^51, 52^. As such, even if granule cell activity exceeds the levels proposed in classical models during behaviors involving diverse cerebellar input, these mechanisms may provide a basis for effective pattern separation.

### Spatio-temporal variability of granule cell representations

Typically, models of cerebellar learning assume consistency of individual granule cell timing across trials, at least for stimulus durations that are appropriate for learning^39^. In contrast, however, our data show that granule cell responses occur with high temporal variability. While previous measurements have shown that the timing of mossy fiber input can be highly reproduceable across trials^53^, this may not translate into a consistency of granule cell spike timing. For example, GABA_B_ receptors^54^, neuromodulators^40, 55^, and cell-intrinsic conductances such as voltage-activated potassium currents can all significantly alter the relationship between synaptic input and granule cell spiking^41^. The reliability of inputs may also be modality specific^56^, as different mossy fibers can have different strengths and short-term plasticity, and likely have different temporal consistency depending on source.

We find that the variability in granule cell spike timing was not dramatically altered when inhibition was blocked. While feedforward inhibition can play a powerful role in establishing spike timing in some circuits^57^, the mossy fiber recruitment of Golgi cell inhibition onto granule cells is relatively weak and inconsistent^58^. Moreover, inhibitory postsynaptic currents onto granule cells often occur before the arrival of excitatory input *in vivo*^59^. Such inhibition evoked before or during excitation likely serves primarily to increase granule cell spike thresholds^13, 60, 61^ rather than as a timing mechanism, consistent with our current observations. As such, refinement of spike timing may not be a primary function of synaptic inhibition in the granule cell layer.

In addition to temporal variability, we observe considerable variability in the identity of granule cell ensembles representing sensory stimuli across trials, an effect primarily due to the relatively low probability of responses on single trials. This low response probability may provide part of the explanation for why cerebellar learning is slow to accumulate across trials, at least as compared to the requirements for induction of synaptic plasticity in the cerebellum when inputs are highly reproduceable^62^.

Despite the spatio-temporal variability in granule cell representations we have measured, it is clear that the cerebellum can form learned associations that are both stimulus-specific and temporally precise^63^. While our measurements support mechanisms of pattern separation, most analogous learning circuits are also thought to involve a secondary process called pattern completion that stabilizes representations. Notably, our observations have been made in the absence of learning, and it remains possible that pattern completion processes in the cerebellar cortex serve to stabilize representations in space and time during learning. Indeed, the cerebellar cortex includes sites of plasticity at almost every node in the circuit, as well as a feedback pathway that provides the type of recurrent structure necessary for pattern completion circuits^64, 65^. Future measurements will be necessary to test how granule cell representations are altered across learning.

### Granule cell inhibition and behavior

We have demonstrated that sensory discrimination in a cerebellum-dependent eyelid conditioning task relies on granule cell inhibition. Specifically, we find that conditioned responses to a CS+ and CS-become indistinguishable when inhibition is blocked. If this change were simply due to mice equally associating the CS+ and CS-, one might expect an increase in the probability of conditioned responses to the CS-. Instead, we observed only a decrease in the response probability to the CS+. This result is consistent with several observations from our imaging data.

First, granule cell tuning was largely preserved when inhibition was blocked, suggesting that the impairment of sensory discrimination is not specifically due to a shift in tuning of the ‘learned’ granule cell population. However, our data also show that blocking inhibition greatly increases the number of responsive cells (Figure 1). Because these emergent cells were not active during conditioning, they were not in the pathway of circuit modifications that occurred during learning. In addition, we find greatly enhanced spontaneous activity across granule cells when inhibition is blocked, an effect that degrades the signal-to-noise ratio of granule cell sensory encoding^61^. We therefore speculate that, together, the enhanced spontaneous activity and the emergent, unlearned CS+ responding cells act to bombard downstream Purkinje cells with non-specific signals that dilute those from the pathways and synapses that were modified during learning. Such results are consistent with a model in which the cerebellum implements a probabilistic binary choice to recognize learned patterns^66^, producing fewer CRs as the learned pattern becomes less discernable during acute block of granule cell inhibition.

Notably, our results partially differ from experiments using large-scale lesions of the cerebellar cortex, which altered both the timing and probability of conditioned eyelid responses in rabbits^67^. The difference in outcome may suggest that the timing mechanism controlling the kinematics of learned eyelid closure is mediated elsewhere in the cerebellar cortical circuit. Along with our observation that granule cell timing is largely inconsistent across trials, our results may suggest that the timing of eyelid responses is primarily computed downstream of granule cells, perhaps at the level of Purkinje cells^68^, where inhibition from molecular layer interneurons directly sculpts the rate and timing of Purkinje cell spiking^69, 70^.

Our data suggest that granule cell representations and behaviors associated with these patterns are highly dependent on inhibitory tone, consistent with findings that chronic hyperexcitability of granule cells can lead to diverse behavioral changes^71^. Because behavior and learning can be context-specific, we suggest that factors regulating granule cell inhibition may provide a powerful basis for enhancing patterns that should be learned or preventing learning in other contexts. In particular, Golgi cells can be bi-directionally modulated by acetylcholine and serotonin^40, 55^. Thus, our measurements suggest that these neuromodulators are likely candidates to enable context-specific cerebellar behavior and learning. These neuromodulators could, for instance, enable the flexibility to express a CR in some contexts but not others.

Together, our results reveal several new mechanisms of cerebellar granule cell layer sensory encoding that depend on local synaptic inhibition. These findings thus significantly extend long-standing predictions of classical Marr-Albus models for how the cerebellar cortex establishes and utilizes discrete sensory representations to guide behavior.

## Methods

### Mice

All experimental procedures using animals were performed with approval of the Duke University Animal Care and Use Committee. Experiments were conducted during the light cycle with both male and female adult mice (>P60). All mice were housed with 12 h light/dark cycles with food and water ad libitum. Imaging experiments were performed with Ai148 (TIT2L-GC6f-ICL-tTA2)-D (Jackson Labs 030328) mice crossed with BACα6Cre-A^21^ (n = 21). Eyelid conditioning experiments used BACα6Cre-A mice (n = 5).

### Surgical procedures

Animals received dexamethasone (3 mg/kg) 3–4 h before surgery. All surgeries were performed under anesthesia, using an initial dose of ketamine/xylazine (50 mg/kg and 5 mg/kg) 5 min before, and 1.0-2.0% isoflurane throughout surgery. Breathing rate and toe pinch responsivity were continuously monitored during surgeries. A heating pad (TC-111 CWE) was used to maintain body temperature. For imaging and eyelid conditioning mice, titanium headplates (HE Parmer) were attached to the skull with Metabond (Parkell). Animals received buprenex (0.05 mg/kg) and cefazolin (50 mg/kg) twice daily for 2 d after surgeries.

For imaging experiments, adult mice (P50–60) were given a 3-mm diameter craniotomy over Crus I at approximately 3.0 mm lateral and 4.3 mm posterior to lambda. Crus I was injected (WPI UMP3) with 150 nL of either AAV7m8-X0117-CAG-DIO-[^+^HTP-GGSGG_8_-GPI-2A-dTomato]-WPRE-pA (HTP_GPI_) (1 × 10^12^; Duke Viral Vector Core) or AAV7m8-6360D-CAG-DIO-[^dd^HTP-GGSGG_8_-GPI-2A-dTomato]-WPRE-pA (^dd^HTP) (1 × 10^12^; Duke Viral Vector Core) at a rate of 30 nl/min and a depth of 350 μm at 2–3 sites. Glass windows consisting of two 3-mm coverslips bonded to a 5-mm coverslip (Warner Instruments No. 1) with index matched adhesive (Norland No. 1) were installed in the craniotomy using Metabond. Imaging mice receiving saline and drug infusions received a plastic cannula (Plastics One; C315GS/PK length 0.5 mm) positioned immediately rostral to the imaging window and attached with Metabond. All mice were given 8 weeks to allow viral expression, including 1–2 weeks of habituation to head restraint.

For eyelid conditioning experiments, adult mice (P50–60) were given 0.3-mm diameter craniotomies at approximately 1.8 mm lateral and 5.85 mm posterior to bregma. Three equidistant 80 nl injections of either AAV-DIO-^+^HTP_GPI_ (1 × 10^12^) or AAV-DIO-^dd^HTP_GPI_ (1 × 10^12^) were performed at a rate of 30 nl/min and a depth of 4.0 mm. Mice receiving saline and drug infusions were implanted with a plastic cannula (Plastics One; C315GS/PK length 3.0 mm) over the injection site that was secured with Metabond. A subset of wild type mice (n = 3) received cannulas but no virus injection at the same location. Mice were given 8 weeks to allow viral expression, including a minimum of 2 weeks for recovery before habituation to head restraints and training.

### Calcium imaging

Two-photon imaging was performed with a resonant scanning microscope (Neurolabware) using a 16X water immersion objective (Nikon CFI175 LWD 16xW 0.8NA). A polymer (MakingCosmetics, 0.4% Carbomer 940) was used to stabilize the immersion solution during imaging. For GCaMP and TdTomato imaging, a Ti:Sapphire laser tuned to 920 nm (Spectra Physics, Mai Tai eHP DeepSee) was raster scanned via a resonant galvanometer (8 kHz, Cambridge Technology) onto the cerebellum at a frame rate of 30 Hz and a field of view of 278 μm × 117 μm (796 × 264 pixels) (Supp. Video 1). Scanbox software (Neurolabware) was used to collect data through a green filter (510 ± 42 nm band filter (SEMrock)) onto GaAsP photomultipliers (H10770B-40, Hamamatsu).

### Behavior

During imaging, animals were head-fixed in a custom sled atop a piezoelectric sensor (C.B. Gitty, 41 mm ‘jumbo’ piezo) read from and triggered through a multifunction data acquisition device (90 Hz, USB X, National Instruments) to measure animal movement^31, 32^. In a subset of experiments, whole-body motion was simultaneously recorded from a CMOS camera (60 Hz, Genie Nano M640 NIR, Teledyne Dalsa) with a fixed focal length lens (6 mm f/2.8, Edmund optics). During imaging, frame-by-frame whisker and facial movements were monitored with the aid of IR LEDs (Swann) from a CMOS camera (60 Hz, Genie Nano M640 NIR, Teledyne Dalsa) with a fixed focal length lens (6 mm f/2.8, Edmund optics) positioned 13 cm above the animal’s head. Piezoelectric and video data were acquired and aligned to imaging data using Scanbox software (Neurolabware) and custom code written in MATLAB (Mathworks). For imaging experiments, sensory stimuli were delivered in pseudorandomized 167 s blocks with randomized inter-trial intervals (ITI) using Mworks (http://mworks-project.org). For somatosensory stimulation, low intensity air puffs (10, 15, or 20 psi, a range of intensities found to produce little or no behavioral response, delivered block-wise) were delivered from a metal tube 5 cm from the center of the vibrissae (5630–10200 ms ITI; 18–29 trials/block). Pure tones of either 68 or 75 dB with frequencies of 1, 5, or 10 kHz were delivered individually block-wise or in randomized pairs for single auditory stimulation (3800–15000 ms ITI; 9–23 trials/block). Imaging sessions lasted between 1.25–2.25 hours.

For eyelid conditioning, the behavioral setup was constructed according to Heiney et al. (2014). Prior to experiments, all mice were habituated to head restraint on the same wheel used for training for 30–60 min/day until they calmly entered head restraints and walked comfortably on the wheel (5–10 d). Stimulus delivery and frame acquisition for video monitoring were triggered with an Arduino Uno microcontroller board (Arduino) controlled with modified Arduino and Matlab code written for Neuroblinks software (Medina lab). Mice were trained during daily sessions of 100–300 trials in which a 50-ms air puff (30 psi) was delivered 3 mm from the mouse’s cornea (unconditioned stimulus, US) and paired with a co-terminating 250-ms, 5 kHz, 70 dB tone (conditioned stimulus, CS). Each session contained one randomly delivered CS only test trial and one US only test trial. Trials were only initiated if the eyelid was open >70–80% for at least 200 ms and at a minimum of 10 s apart^43^.

### In vivo pharmacological infusions

Saline and drugs were infused into awake, head-fixed mice using an automated pump (WPI UMP3), a Hamilton syringe (10 μl Gastight model 1701 RN) and a plastic internal cannula (Plastics One, C315IS) threaded into the guide cannula. All infusions had a total volume of 1 μl delivered at a rate of 1 μl/min. To estimate the spread of pharmacological agents under the imaging window, 10 mM fluorescein dye (Sigma-Aldrich #F6377) dissolved in sterile artificial cerebrospinal fluid (aCSF; 150 nM NaCl, 4 mM KCl, 2 mM MgCl2, 2 mM CaCl2, 10 mM HEPES, 10 mM glucose, pH 7.4) was infused through the cannula rostral to the imaging window, followed by a 1 h rest and perfusion. For all other imaging and behavior experiments, aCSF only or either 1 μM non-binding gabazine.1^nbDART^ or 1 μM gabazine.1^nbDART.2^ was dissolved in sterile aCSF and applied, followed by a 20-min rest. Infusions were delivered at least 1 d apart for each animal. At least one day after gabazine.1^nbDART^ administration, 1 μM Alexa647.1^DART.2^ was dissolved in sterile aCSF and infused, followed by 1 h rest and perfusion. In a subset of eyelid conditioning experiments, fluorescent muscimol (1 mM; BODIPY TMR-X muscimol conjugate; Invitrogen) was infused in wild type mice, followed by a 3-min rest period.

### Histology

Mice were anaesthetized with an IP injection of ketamine/xylazine (200 mg/kg and 30 mg/kg, respectively) prior to perfusion with PBS and 4% paraformaldehyde. 50 μm sagittal sections were cut using a vibrotome (Pelco 102). Slices were mounted using Southern Biotech DAPI-Fluoromount G or Vectashield Vibrance (Vector labs) then imaged using an upright confocal microscope (Leica SP8).

### Slice Electrophysiology

Acute brain slices and associated whole-cell electrophysiological recordings of synaptic inhibition were performed as described previously^i40^.

### Data Analysis

#### Multi-photon imaging

All acquired two-photon images were processed using a combination of the opensource Python toolbox for large scale calcium imaging data analysis CaImAn^72^ and custom code written in MATLAB. First, images were corrected for motion over 60 × 60-pixel patches using a piecewise rigid motion correction algorithm (NoRMCorre;^73^). All videos were manually screened to ensure adequate motion correction. Whole experiments that could not be made to produce a stable averaged image were excluded from further analysis. Images collected before and after drug infusions, without displacing the objective, were motion registered and segmented together to enable reliable comparison between conditions. Then, source separation was performed using constrained non-negative matrix factorization (CNMF; ^74^). This algorithm includes exclusion of fluorescence changes originating in the neuropil. Regions of interest (ROI) identified by CNMF were then sorted according to spatial stability, transient signal-to-noise ratio, and performance in a CNN based classifier^72^. ROIs were then excluded based on their proximity to the edge of the FOV and overlap with non-specifically-labelled structures (i.e., anything other than putative granule cells) in motion-registered, averaged images using custom MATLAB code. Remaining raw Ca2+ time courses computed by CNMF were screened for periods in which the signal exceeded 6 standard deviations from the mean of either the first or last 20,000 imaging frames for >2 s, as such changes were noted to occur in cells that become bright and swell over the course of an experiment and were presumed to be unhealthy/dying. ROIs with this fluorescence signature and a bright, swollen appearance in motion-registered, averaged images were excluded from further analysis. Additionally, raw Ca2+ time courses lacking stability by 1) slowly drifting in magnitude or 2) transiently or permanently losing all signal were excluded from further analysis to allow reliable comparison of responses throughout each experiment. Fluorescence changes (ΔF) were normalized to a 1-s window of baseline fluorescence prior to stimulus onset for each trial. Individual cell responses were considered significant if they surpassed 2 standard deviations from the baseline period between 90–180 ms after stimulus onset (somatosensory stimulation), or if any sliding window beginning at stimulus onset and ending 0.1 s after stimulus offset surpassed 2 standard deviations of any equal-length window during the baseline period (auditory stimulation). Some cells with significant responses in the control condition no longer achieved significance after DART infusions due to high levels of activity in the baseline period, despite having activity during the stimulus windows. Because this reduced signal-to-noise impaired accurate measurement of sensory responses, cells that lost significance in DART were not included for condition-matched analyses. The fraction of responsive granule cells for each condition was estimated by calculating the number of granule cell sized ROIs (∼14.7 pixel diameter) that would tile the FOV (796 × 264 pixels) without overlap.

First and peak event times and amplitudes during individual trials were calculated using trapezoidal numerical integration, identifying peaks ≥120 ms. ‘Off’ cells were defined as cells having a mean first peak time at or after stimulus offset.

### Unisenosry stimulus classification

To classify auditory stimuli, calcium signals from populations of granule cells were used. First, noise was removed in two steps. 1) Smoothing: calcium signals were first smoothed by a 3-frame boxcar filter. 2) Threshold: after smoothing, baseline noise was estimated from 25 frames (30 Hz sampling) preceding the stimulus. Events that exceeded 2-SD above the baseline noise were retained, signals below this threshold were set to zero. A granule cell response was defined as the peak calcium signal (between the initiation of the sound and 5 frames after it ended) on each trial after smoothing and thresholding. A population response was defined by accumulating this peak calcium signal across all granule cells for a given trial. This created a matrix of responses that was [cells x trials]. Given a limited number of trials in many experiments (∼20) and a large population response (∼300 cells), response classification was performed using a non-parametric nearest-neighbor approach^75^. This approach computed the distance of a test trial to all training trials. The test trial was classified according to the stimulus condition that produced the nearest response among the training trials. Test-trials were selected from a ‘hold-one-out’ approach and the remainder of the data was used for training. Three classification tasks were run: identify the identify the frequency of the stimulus (three categories, Fig. 4e left); identify the amplitude of the stimulus (two categories, Fig. 4e middle); and identify the amplitude and frequency of the stimulus (six categories, Fig. 4e right). Classification was performed across all trials. For stimulus classification under control conditions, responses from 328 granule cells were used with 23 trials for each condition. For the gabazine.1^DART.2^ discrimination, 319 granule cells were used with 22 trials for each condition.

Several control analyses were run to ensure the discrimination results were not overly sensitive to changes in the procedure described above. First, half the number of cells were tested in each condition, to make sure the results were relatively insensitive to this parameter. The results in Figure 4 were qualitatively similar; none of the trends or statistically significant differences changed. Second, we defined responses in several different ways. In addition to using the peak amplitude (described above), we also used: 1) the time of the peak response; 2) the time and amplitude of the peak response; 3) the time the response initially crossed 2-SD above baseline; 4) the time and amplitude the response initially crossed baseline; 5) the integrated calcium signal between frame 30 (when tone was initiated) and 65 (five frames after it ceased); 6) the integrated calcium signal from ‘5)’ and the initial time it crossed the significance threshold. The results in Figure 4 were qualitatively similar; none of the trends changed, but under some response definitions, some differences failed to clearly reject the null hypothesis (p-values were > 0.05). In Figure 4, error bars represent SEM and were computed by using the Wald method of confidence interval estimation for binomial distributions.

The same procedures were used to classify puff stimuli. Responses were classified using a population of 275 granule cells with 15 trials for each stimulus amplitude under control and DART conditions. Decoding was performed on the peak response between frames 27 and 37 on each trial.

### Unisensory versus multisensory response discrimination

Actual multisensory (sound and puff) responses were not the linear combination of unisensory (sound or puff) responses. To examine the consequences of this deviation from linear signal summation on stimulus discrimination (multi-versus unisensory), we simulated linear multisensory responses by summing unisensory responses sampled from responses to either auditory stimulus with responses to the puff stimulus for each cell. Discrimination performance (correctly discriminating between unisensory and multisensory responses) was then measured using multisensory responses or the simulated linear multisensory responses (Fig. 5). Response classification was performed using a non-parametric nearest-neighbor approach^75^. Responses were derived from 521 granule cells and 30 multisensory and 30 unisensory trials (10 auditory trials at two frequencies and 10 puff). Calcium signals were smoothed and thresholded as described in the previous subsection. Responses were defined as the integrated calcium signal starting at the time the signal crossed threshold after the initiation of the auditory stimulus until 5 frames (30 Hz sampling) after the termination of the auditory stimulus. Qualitative results and significance tests were robust to halving the number of cells and/or using other definitions of ‘response,’ such as the amplitude of the peak response and the time-of-peak response.

### Behavior Analysis

For imaging experiments, a machine-learning-based algorithm (DeepLabCut^76^) was used to automatically track components of the face, whiskers, and head in accompanying high-speed videos. Tracked features were initially labeled manually in a small portion of frames (30) to train the algorithm, then x and y locations of each feature were automatically determined for all remaining frames. Motion was evaluated as cumulative displacement of these coordinates during aligned calcium imaging frames.

To validate the effectiveness of the piezo sensor at detecting motion, the same sensory stimuli used in imaging experiments were delivered while collecting high-speed video of each mouse’s head and limbs. Machine learning-based motion tracking with DeepLabCut^30^ revealed that the sensor reliably detects limb motion and other movements such as grooming (Supp. Fig. 5). Specifically, limb and facial movements were aligned with piezo traces, revealing that piezo measurements reliably reflect movements of all four limbs, as well as fine movements of the ears and face. Video detected motion and piezo recordings do not have a one-to-one relationship, however, and some changes in piezo voltage do not correspond with any visible movement detected by video. We interpret these changes in piezo voltage as likely to reflect muscle tension as the mouse prepares to move, as they generally occur immediately preceding video detected movements. However, nearly all video-detected movements are less than 5 frames away from piezo deflections that are >1 standard deviation from the mean, so this threshold was used to segregate imaging frames recording during movement. Trials were excluded from analysis if movement occurred anytime between the second before stimulus onset and 300 ms after stimulus offset.

In addition to removing signals related to movement, piezo voltage traces identify frames in which significant animal movement causes failures of imaging motion correction. Two-color imaging of both neural activity with GCaMP6f and tdTomato indicates that instances of tdTomato fluorescence fluctuations (indicating Z-motion or another motion correction failure) are also excluded from analysis using the above criteria (Supp. Fig. 5).

Because mice whisk frequently, and granule cells in Crus I can be tuned to whisker movements^25, 27^, we used a different approach to segregate cells that are modulated during whisking. During imaging, videos of the head and whiskers were used to align whisker movements to changes in granule cell activity. Because some granule cells can be tuned for whisker deflection angle^25^, whisker movements were then parsed according to movement amplitude, and whisks that occurred between trials and in the absence of whole-body movement were used to identify granule cells that were modulated specifically by whisking. Whisk modulated cells overlapped very modestly with ensembles responsive to auditory or somatosensory stimulation (7.7 ± 4.5% of sensory and whisking responsive cells). Accordingly, inclusion of whisk modulated cells had no significant effect on auditory (unpaired t-test, *P*=0.9686) and somatosensory (unpaired t-test, *P*=0.4864) responses on average. (Supp. Fig 4).

For eyelid conditioning experiments, behavioral data were analyzed using modified Matlab code written for Neuroblinks software (Medina lab). Briefly, fraction of eyelid closure was calculated for each video frame by generating a binary image of a region of interest surrounding the eye, thresholded to provide maximal discriminability for each experiment, then summing pixel counts for each frame. Conditioned responses (CRs) were defined by eyelid closure >10%. CR probability for a given session was calculated according to all paired trials during that session. CR amplitudes were calculated as the mean closure during a 4-frame window preceding the US.

### Study Design

Sample sizes were similar to sample sizes used for comparable studies in the field: 3 mice or more per condition were used for imaging and eyelid conditioning experiments, with the exception of only 2 mice used for ddHTP imaging experiments. Each imaging experiment included over 150 active cells. No statistical methods were used to select sample sizes. Data exclusions and rationale are indicated in relevant sections above. All major results were replicated in multiple mice. Additionally, major results were replicated in mice with viral, rather than transgenic, calcium indicator expression and using different analytical approaches. Mice were randomly selected for experimental groups. Trial types were pseudo-randomized during imaging experiments and randomized by behavioral software for eyelid conditioning experiments. Investigators were not blinded to mouse group during experiments or analysis.

### Data and Code Availability

Data and code that support the findings of this study are available from the corresponding authors upon reasonable request.

## Acknowledgements

This work was supported by grants from the NIH NINDS (CH: 5R01NS096289-02, EF: 1F31NS113742-01A1). We thank Drs. Stephen Lisberger, Lindsey Glickfeld, John Pearson, and Nicole Calakos for input throughout the project, Drs. Javier Medina and Shane Heiney for input specific to eyelid conditioning experiments, Drs. Javier Medina and Wade Regehr for comments on the manuscript, and Dr. Brenda Shields for technical assistance with DART reagents.

## Figure Legends

**Supplemental Figure 1.**
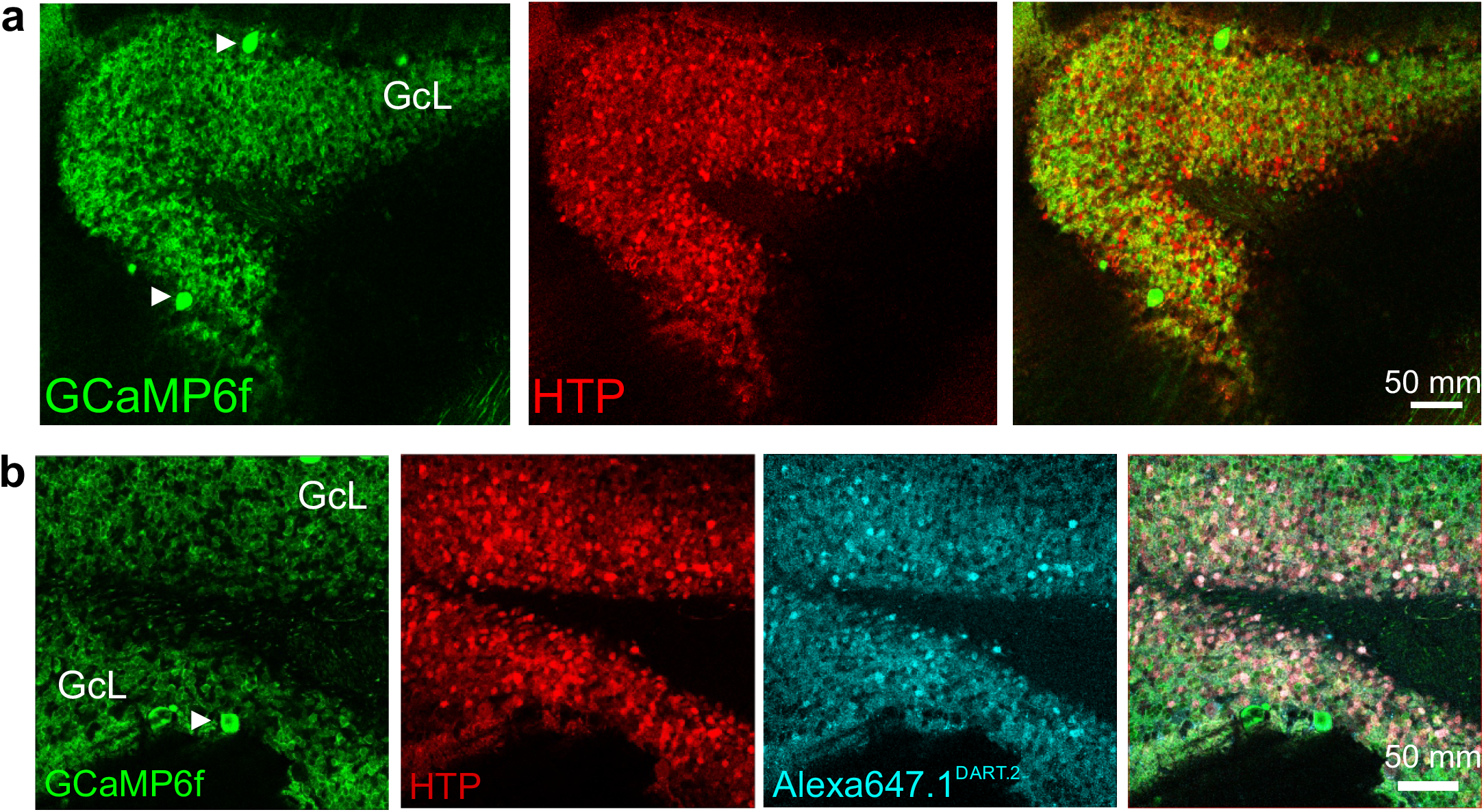
Selective labeling of cerebellar granule cells. **a**. Example confocal images from a representative mouse expressing GCaMP6f (left) and HTP (middle) and their intersection (right) in the granule cell layer (GcL). Arrows note sparse labeling of Purkinje cells, which were excluded from analysis. **b**. Same as **a**, for a different representative mouse infused with Alexa647.1^DART.2^ (cyan, middle right) and intersection of all three labels (right).

**Supplemental Figure 2.**
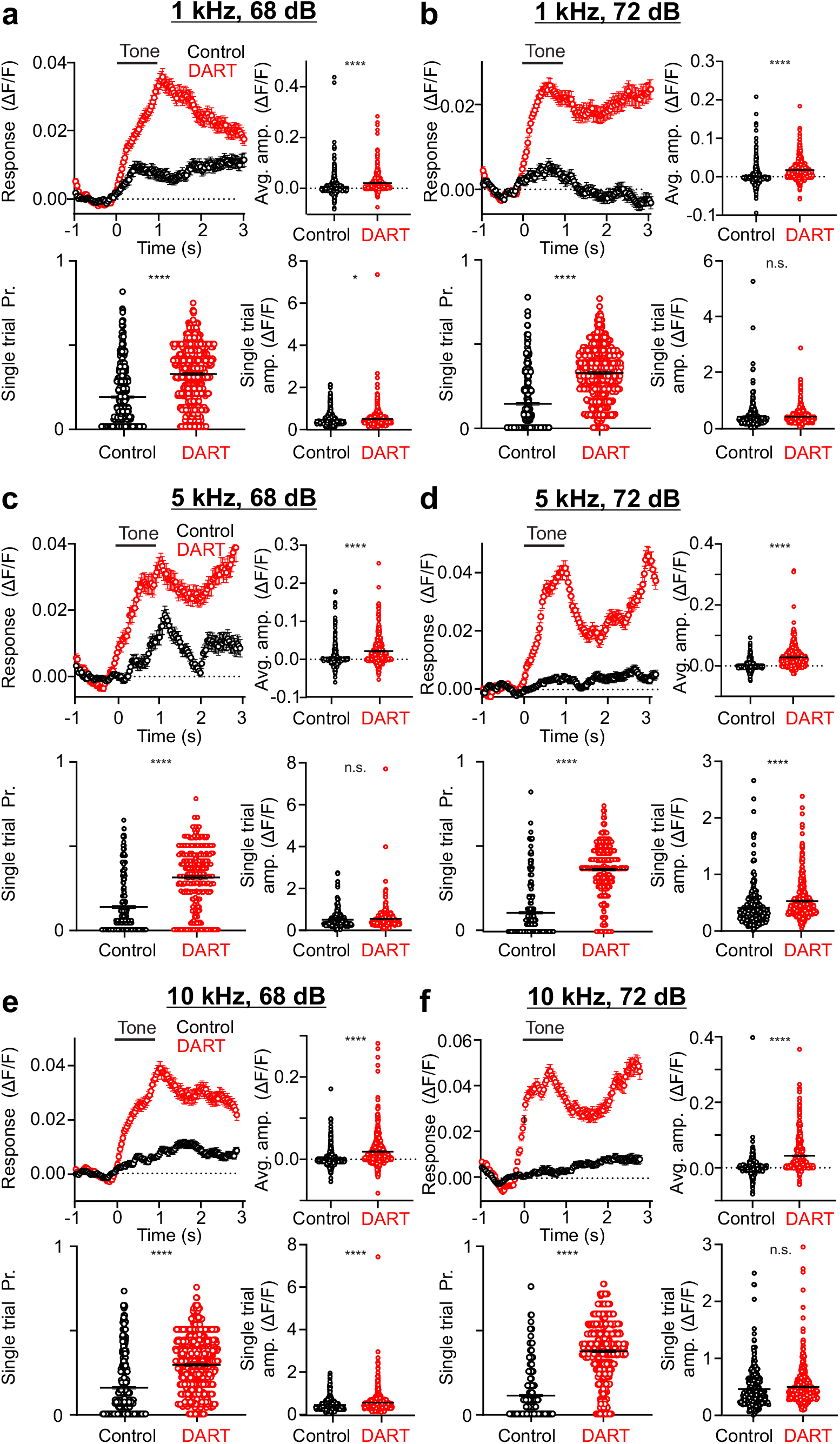
Local synaptic inhibition sparsens and thresholds cerebellar granule cell responses to all auditory stimuli presented. **a**. Top left, mean time course of responses during 1 kHz, 68 dB tone presentation before (black) and after (red) DART infusion. Error is SEM across cells. Top right, mean response amplitudes for individual cells before (black) and after (red) DART infusion. Black lines are mean ± SEM across cells. Bottom left, same as top right, for response probability. Bottom right, same as top right, for mean responses on all trials with significant responses (n=689 cells, n=458 cells for single values; 6 mice). **b**. Same as **a**, for 1 kHz, 72 dB tones (n=541, n=296 for single trial values; 5 mice). **c**. Same as **a**, for 5 kHz, 68 dB tones (n=381; n=184 for single trial values; 3 mice. **d**. Same as **a**, for 5 kHz, 72 dB tones (n=493cells, n=251 for single trial values; 3 mice). **e**. Same as **a**, for 10 kHz, 68 dB tones (n=638 cells, n=409 cells for single trial values; n=6 mice). **f**. Same as **a**, for 10 kHz, 72 dB tones (n=566; n=292 for single trial values; 3 mice). \**p* < 0.05, \*\*\*\**p* < 0.0001.

**Supplemental Figure 3.**
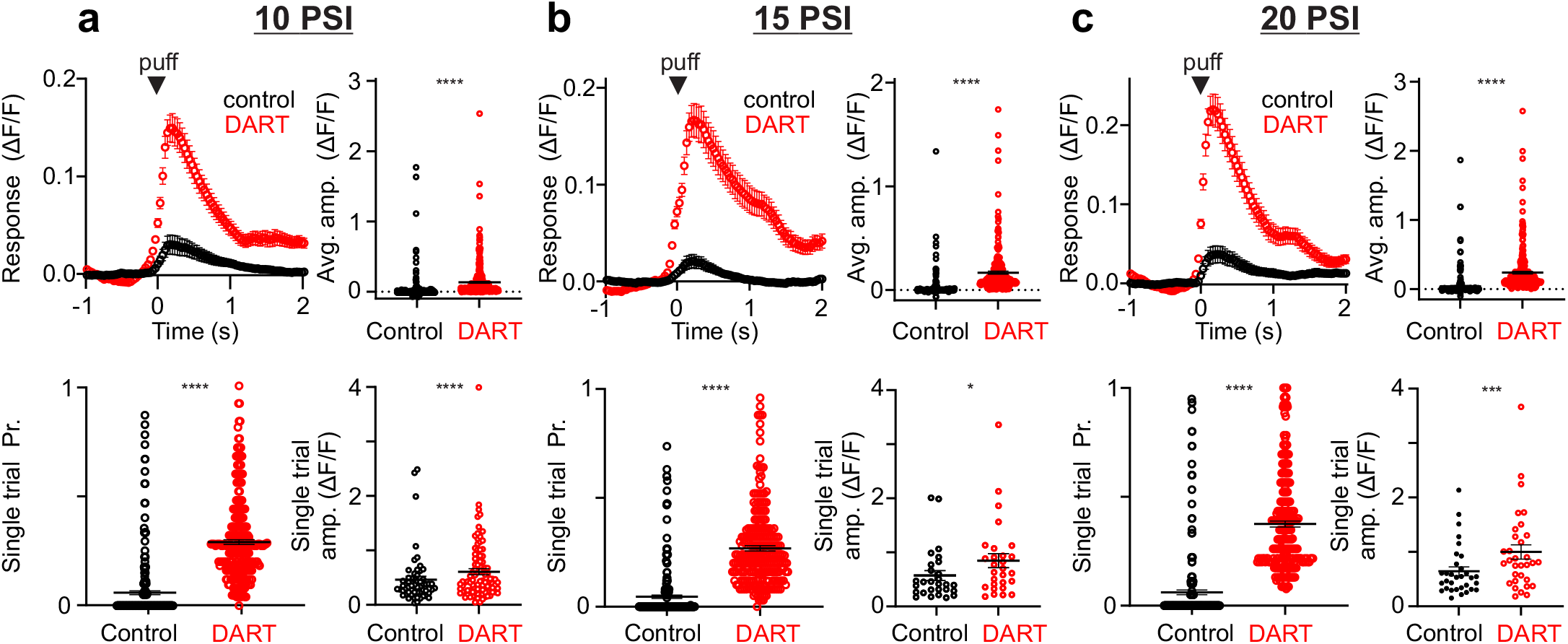
Local synaptic inhibition sparsens and thresholds cerebellar granule cell responses to all somatosensory stimuli presented. **a**. Top left, mean time course of responses to a 10 PSI air puff before (black) and after (red) DART infusion. Error is SEM across cells. Top right, mean response amplitudes for individual cells before (black) and after (red) DART infusion. Black lines are mean ± SEM across cells. Bottom left, same as top right for response probability. Bottom right, same as top right for mean responses on all trials with significant responses (n=283 cells; n=58 cells for single trial values; 6 mice). **b**. Same as **a**, for 20 PSI air puff (n=234, n=28 for single trial values; 3 mice). **c**. Same as **a**, for 30 PSI air puff (n=261, n=32 for single trial values; 3 mice).

**Supplemental Figure 4.**
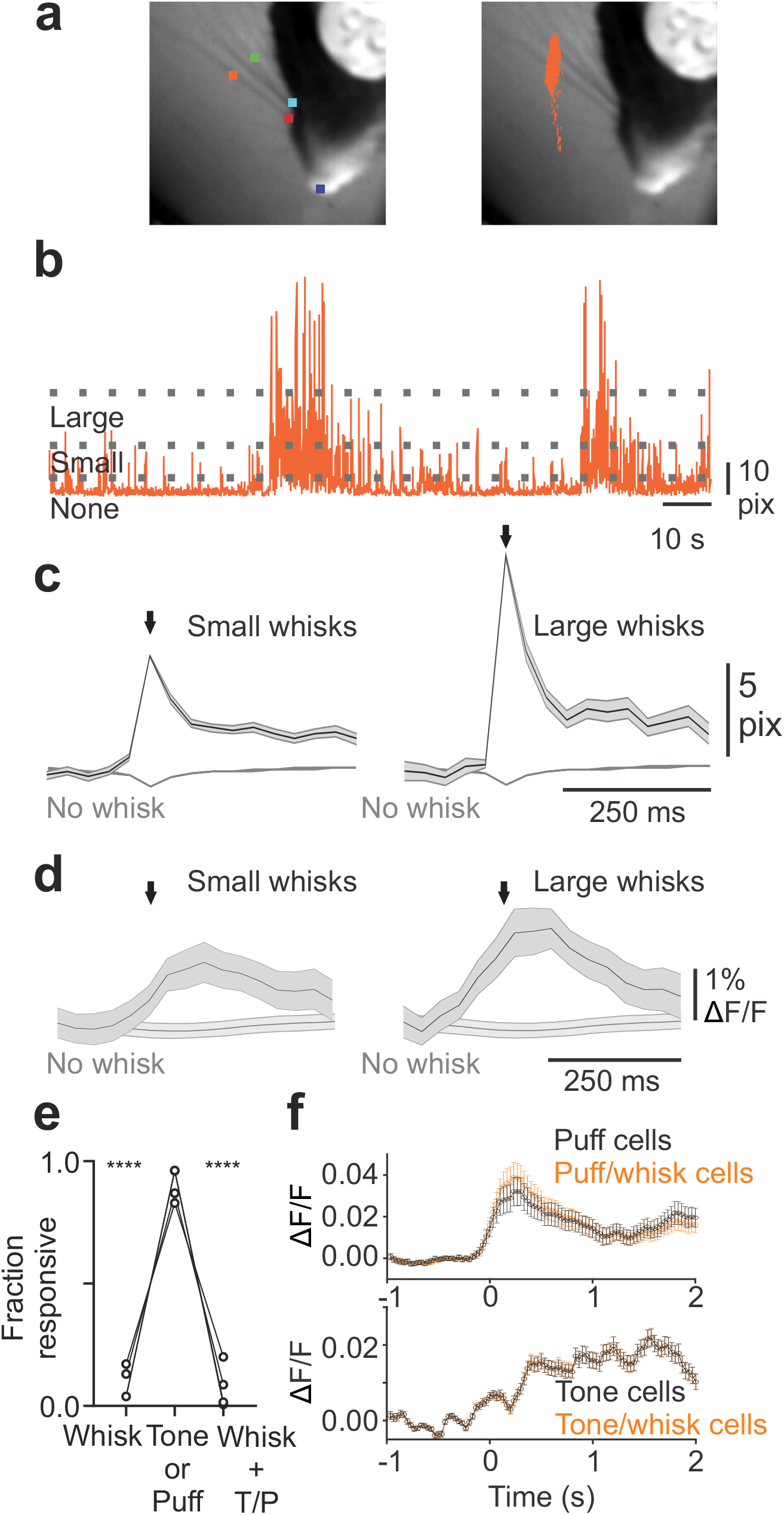
Tracking of whisker movements with high-speed video during imaging enabled identification of whisking modulated cells and evaluation of their contribution to sensory responses. **a**. Left, example image from a representative mouse with features labeled for automated identification of two whisker root (light blue and red) and midpoints (orange and green), as well as a stationary feature (snout, dark blue) used to estimate tracking jitter. Right, image from **a** with x and y coordinates for anterior whisker midpoint for all frames overlaid. **b**. Time course of displacement (in pixels) of whisker midpoint labeled in **a**. Dashed lines indicate thresholds used to segregate whisks according to magnitude. **c**. Left, mean ± SEM of low amplitude whisks occurring between trials and in the absence of body movement during an example experiment (n = 816 whisks). Mean ± SEM of frames without whisker movement plotted behind. Right, same for high amplitude whisks measured during the same experiment (n = 287 whisks). **d**. Left, mean ± SEM fluorescence of cells modulated during small whisks and fluorescence of the same cells during frames without movement (n = 16). Right, same for independently identified cells modulated during large whisks (n = 16). Cells had the same identity. **e**. Fraction of responsive cells from 4 mice with significant activity during whisking, a sensory stimulus (auditory or somatosensory), or both; RM ANOVA: p = 0.0013, Tukey’s multiple comparison’s test: stim v. whisk: p=0.0025, stim v. w+s: p = 0.0036, whisk v. w+s: *P* = 0.5782). **f**. Top, Time course of mean responses to a somatosensory stimulus for all cells significantly responsive to the stimulus (orange; n = 366) and cells significantly responsive to the stimulus not modulated during whisking (black; n = 306; paired t-test of mean during stimulus window: control = 0.0343 ± 0.0066, DART = 0.0278 ± 0.0065, *P* = 0.4864). Error is SEM. Bottom, same as top for cells significantly responsive to an auditory stimulus (orange: n = 432; black: n = 424; paired t-test of mean during stimulus window: control = 0.0123 ± 0.0015, DART = 0.0122 ± 0.0014, *P* = 0.9686).

**Supplemental Figure 5.**
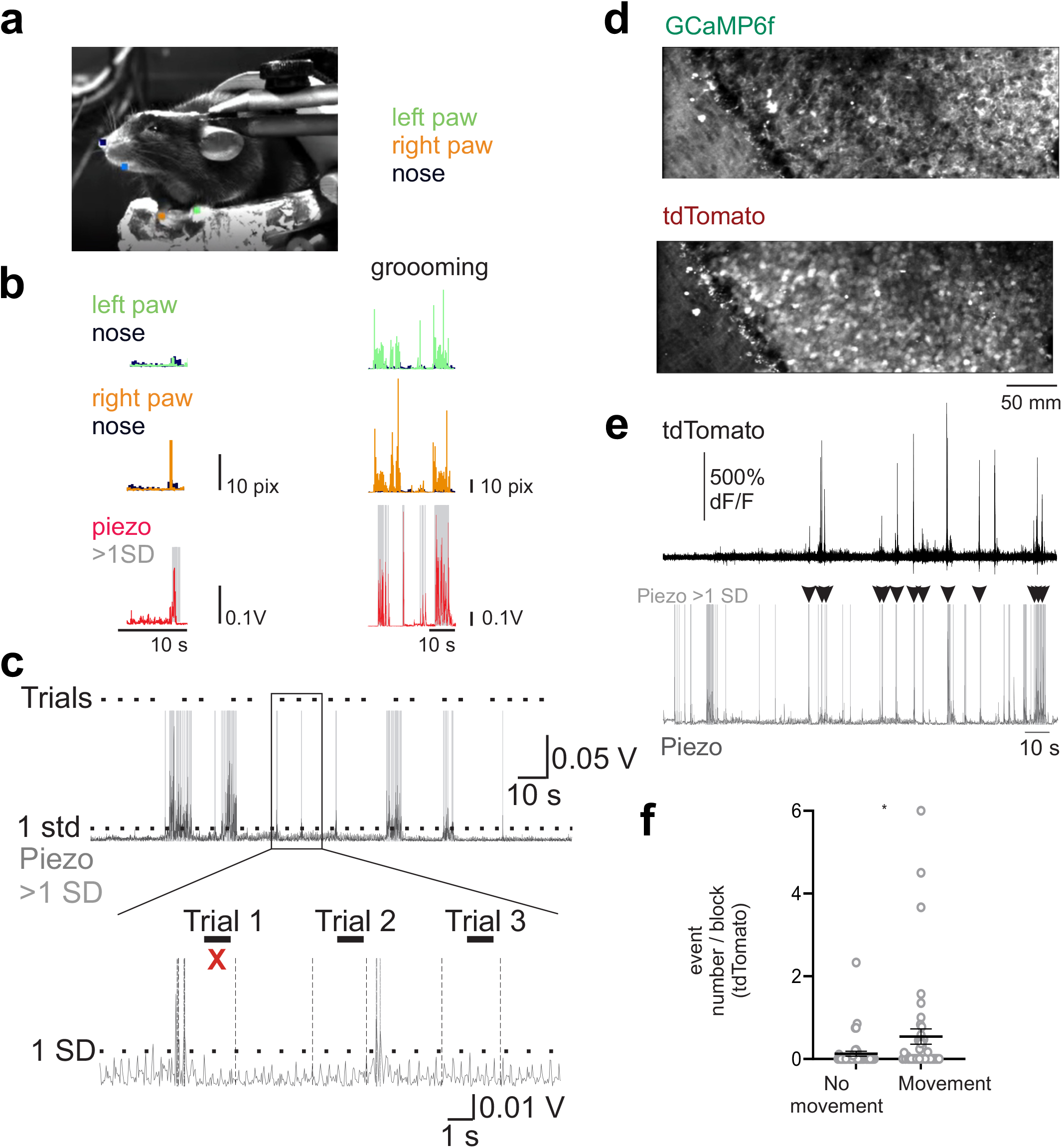
Piezo vibration sensor reports body movements and facilitates removal of trials with z-motion. **a**. Example image from a representative mouse following automated identification of left (green) and right (orange) paws, nose (dark blue) and mouth (light blue). **b**. Time course of displacement (in pixels) of left paw and nose (top), right paw and nose (middle) during left paw lift (left) and grooming (right). Bottom, simultaneous motion measurement using a piezoelectric sensor under the mouse’s torso. Shaded gray regions reflect motions greater than 1 standard deviation that were discarded. **c**. Example raw piezo sensor trace at low (top) and high (bottom) temporal resolution. Dashes at top of each trace indicate times of auditory or somatosensory stimulus presentation; dashed line indicates 1 standard deviation threshold. Shaded regions indicate epochs that exceed this threshold; red x indicates trial that was discarded. **d**. Example field of view with GCaMP6f (top) and tdTomato (bottom) expression. **e**. Time course of changes in fluorescence in the tdTomato channel (top) after x-y motion correction and fluctuations in the piezo sensor. Note that changes in tdTomato fluorescence correspond with periods of animal movement (arrowheads), likely corresponding to uncorrected effects of z-motion. **f**. Quantification of the mean number of tdTomato events detected during blocks in which the mouse was stationary or moving (n = 44 blocks; paired t-test P = 0.00177). Error is SEM across blocks.

**Supplemental Figure 6.**
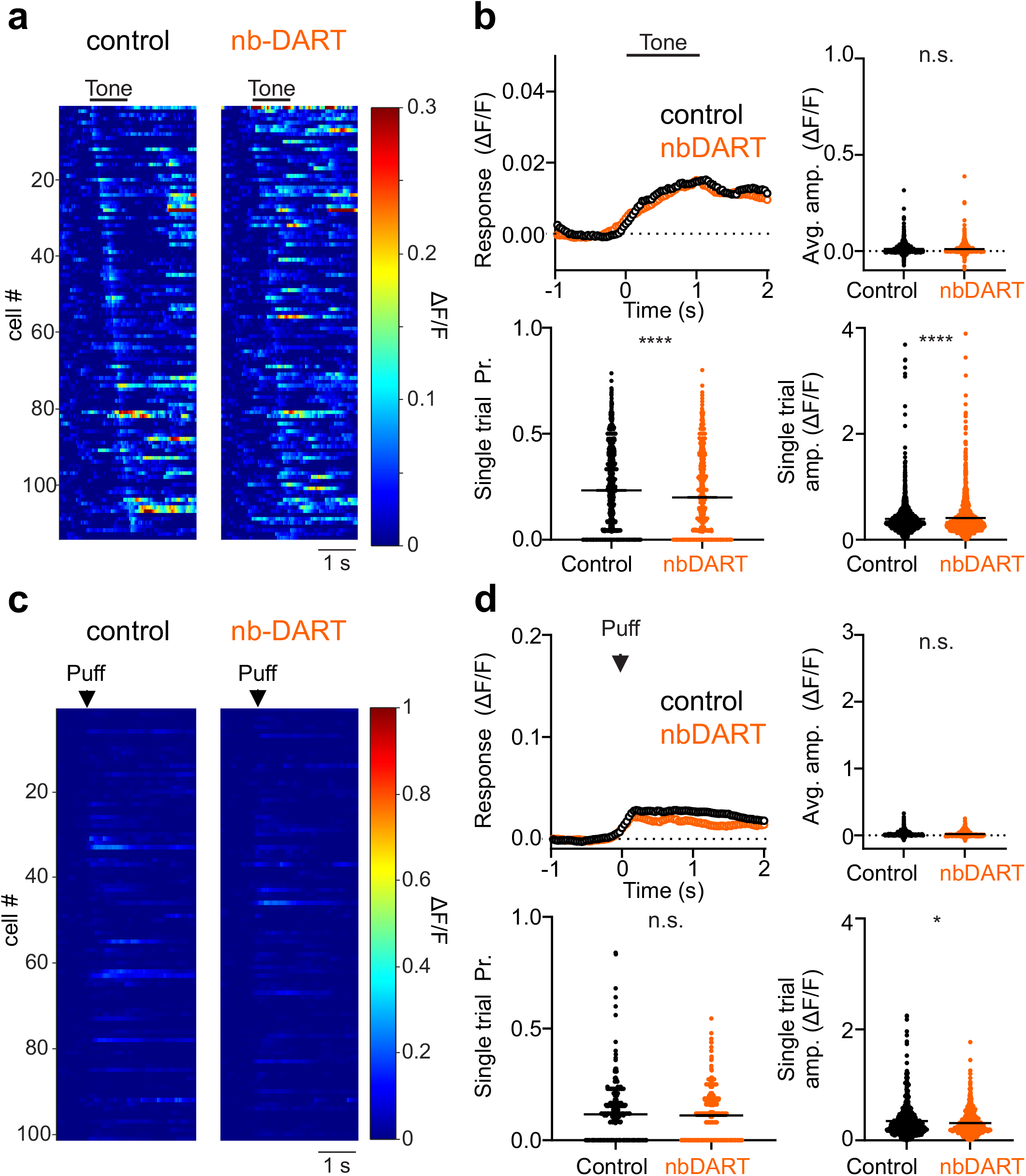
Probability and average amplitudes of auditory and somatosensory responses not changed after infusion of gabazine.1^nbDART^ variant. **a**. Heatmap of average responses to auditory stimuli in control conditions (left) and after infusion of **gabazine.1**^**nbDART**^ (nbDART, right) in an example mouse with HTP expression. **b**. Top left, mean time course of responses during tone presentation before (black) and after (orange) nbDART infusion. Error is SEM across cells. Top right, mean response amplitudes for individual cells before (black) and after (orange) nbDART infusion. Black lines are mean ± SEM across cells. Bottom left, same as top right, for response probability. Bottom right, same as top right, for mean responses on all trials with significant responses (n = 2044, n = 1114 for single trial values; 3 mice). **c-d**. Same as **a-b**, for responses to somatosensory air puffs (n = 770, n = 259 for single trial values; 3 mice). \**p* < 0.05, \*\*\*\**p* < 0.0001.

**Supplemental Figure 7.**
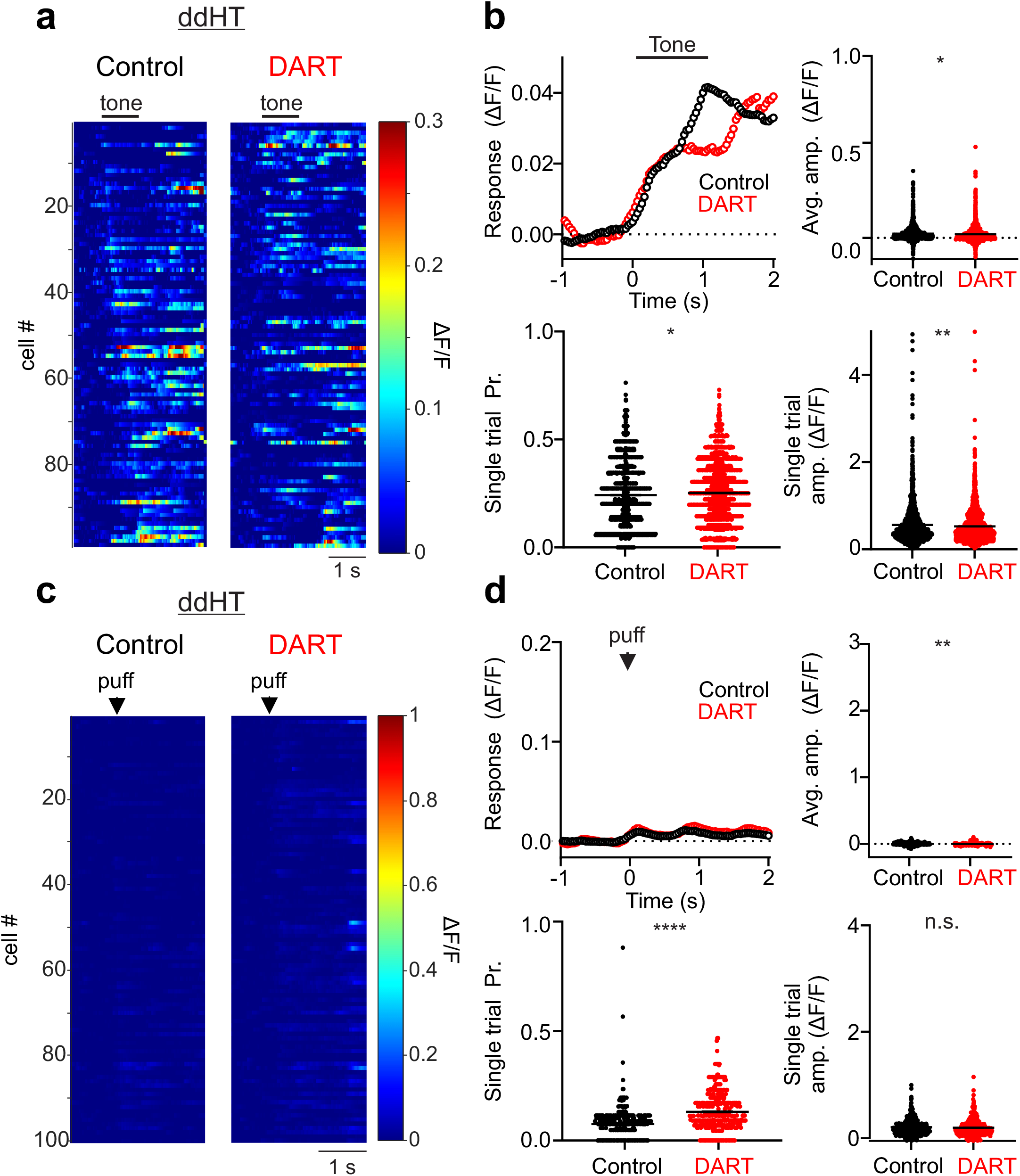
Probability and average amplitudes of auditory and somatosensory responses not changed after infusion of DART with expressed non-binding ^dd^HTP. **a**. Heatmap of average responses to auditory stimuli in control conditions (left) and after infusion of DART (right) in an example mouse with ^dd^HTP expression. **b**. Top left, mean time course of responses during tone presentation before (black) and after (red) DART infusion. Error is SEM across cells. Top right, mean response amplitudes for individual cells before (black) and after (red) DART infusion. Black lines are mean ± SEM across cells. Bottom left, same as top right, for response probability. Bottom right, same as top right, for mean responses on all trials with significant responses (n = 1853, n = 1723 for single trial values; 2 mice). **c-d**. Same as **a-b**, for responses to somatosensory air puffs (n = 443, n = 266 for single trial values; 2 mice). \**p* < 0.05, \*\**p* < 0.01, \*\*\*\**p* < 0.0001.

**Supplemental Figure 8.**
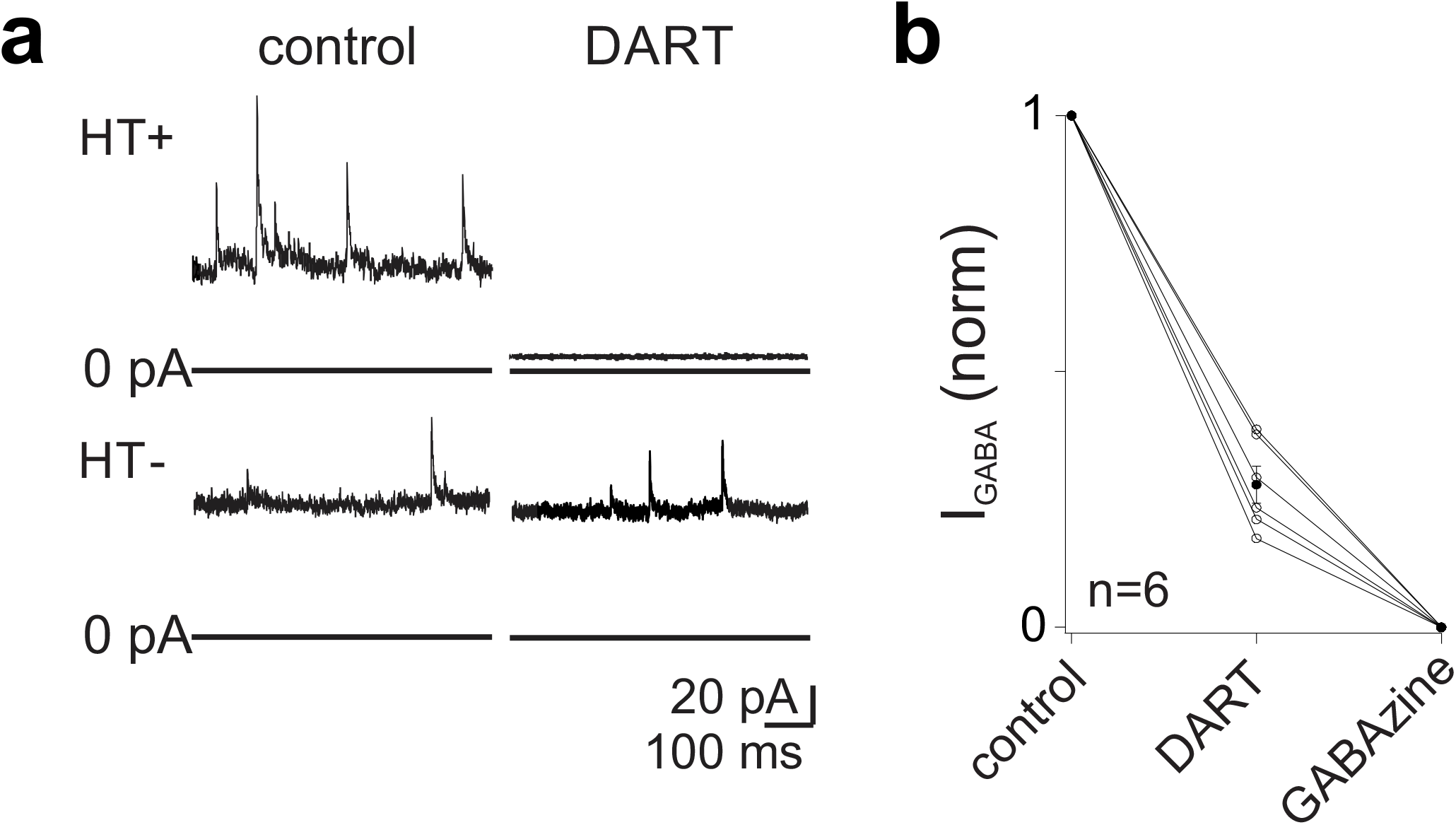
gabazine.1^DART.2^ blocks GABAergic inhibition onto granule cells *in vitro*. **a**. Top, Example voltage-clamp recording at 0 mV from a granule cell expressing HTP. Wash-in of 100 nM gabazine.1^DART.2^ (10x lower than *in vivo* infusions) blocks GABAergic inhibition, both spontaneous IPSCs and the tonic component of GABAergic inhibition. Bottom, the same concentration of DART does not block inhibition in a granule cell that does not express HTP. **b**. Across HTP expressing cells, 100 nM gabazine.1^DART.2^ blocked the total GABAergic current (sIPSCs + tonic current) by 73 ± 4%. All procedures for *in vitro* voltage-clamp measurements from granule cells were performed as described previously^40^.

**Supplemental Figure 9.**
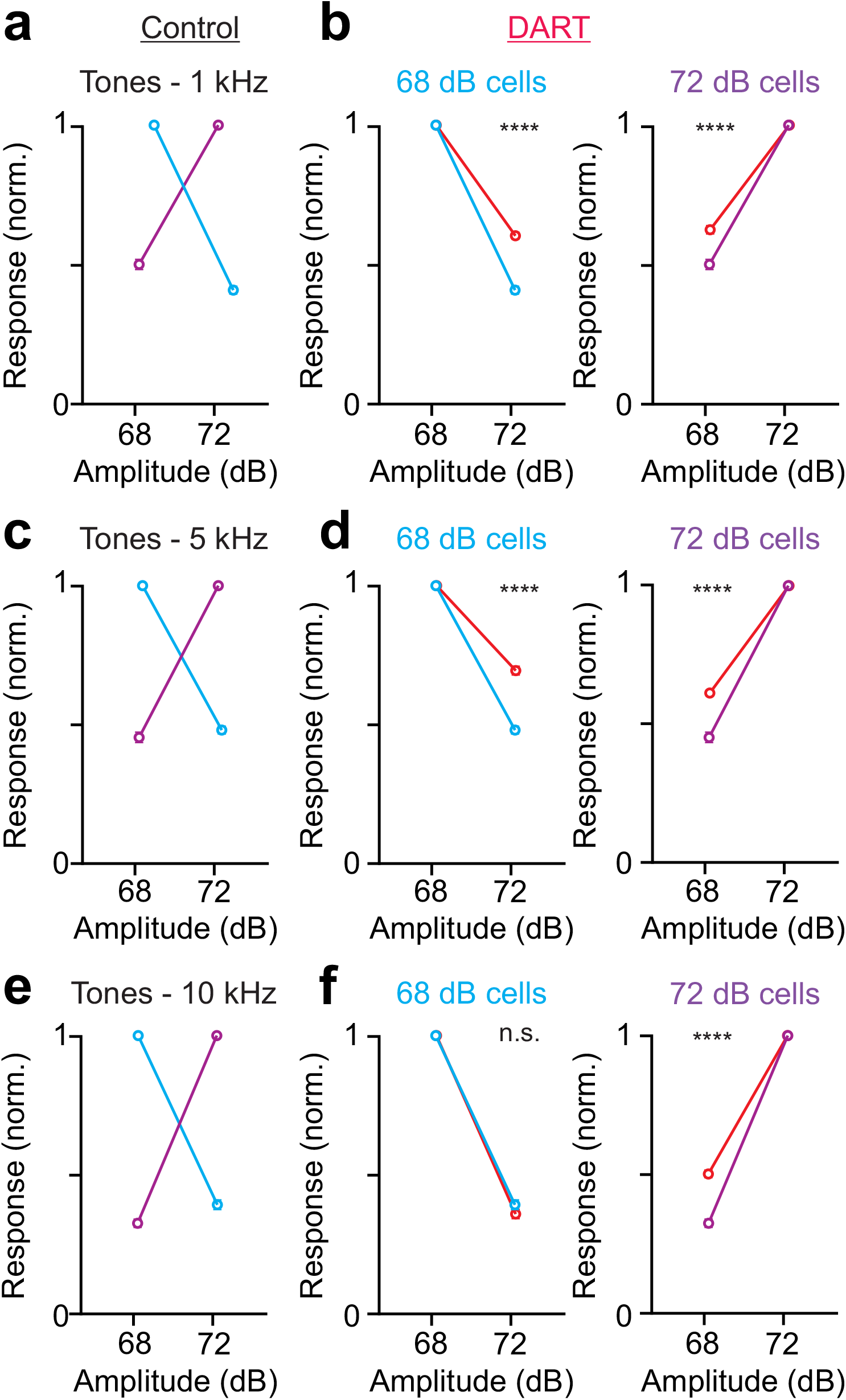
Granule cells display preferences for auditory stimulus amplitude. **a**. Normalized responses for cells that prefer 68 dB (cyan; n = 342) or 72 dB (purple; n = 207) in response to 1 kHz tones. Error is SEM across cells. **b**. Normalized responses for cells that prefer 68 dB (left) or 72 dB (right) in control (cyan/purple) and after DART infusion (red; 68 dB: n = 390), 72 dB: n = 323). Data is the same as in **a** for control conditions. **c-d**. Same as **a-b**, for 5 kHz tones: control, 68 dB: n = 293, 72 dB: n = 213; DART, 68 dB: n = 220, 72 dB: n = 469 **e-f**. Same as **a-b**, for 10 kHz tones: control, 68 dB: n = 241, 72 dB: n = 367; DART, 68 dB: n = 381, 72 dB: n = 344.

**Supplemental Figure 10.**
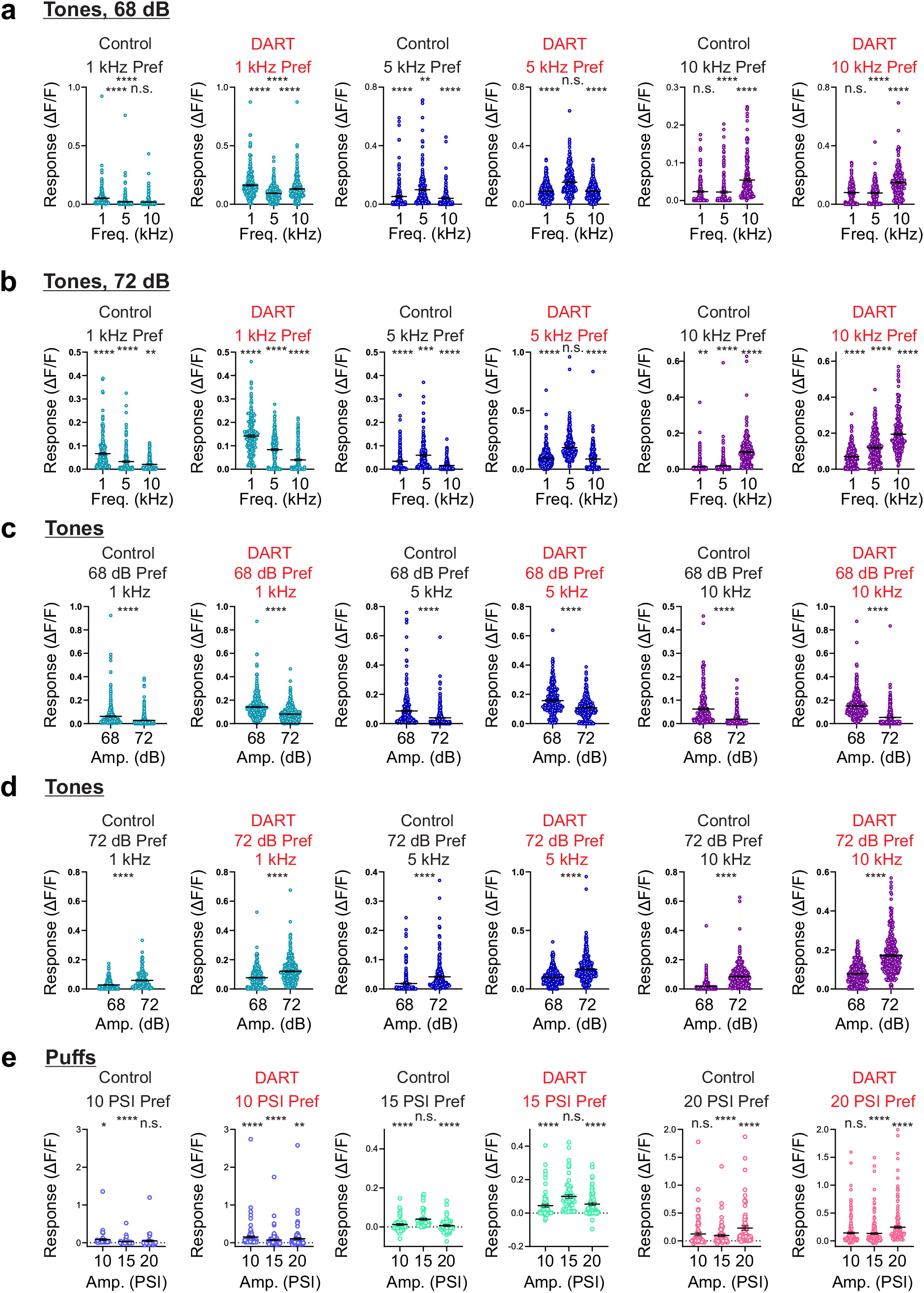
Granule cells display preferences for auditory stimulus frequency and amplitude and somatosensory stimulus amplitude before and after blocking synaptic inhibition. **a**. Left, maximum responses (ΔF/F) for cells significantly responsive to any 68 dB tones that prefer 1 kHz in control (n = 227) and DART (n = 250) conditions. Middle, same as left for cells that prefer 5 kHz: control: n = 211; DART: n = 297. Right, same as left for cells that prefer 10 kHz: control: n = 166; DART: n = 169. **b**. Same as a for cells significantly responsive to any 72 dB tones: 1 kHz, control: n = 184, DART: n = 170; 5 kHz, control: n = 141, DART = 331; 10 kHz, control: n = 313, DART: n = 224). **c**. Left, maximum responses (ΔF/F) for cells significantly responsive to 1 kHz at any amplitude that prefer 68 dB in control (n = 342) and DART (n = 207) conditions. Middle, same as left for cells responsive to 5 kHz that prefer 68 dB in control (n = 293) and DART (n = 220) conditions. Right, same as left for cells responsive to 10 kHz that prefer 68 dB in control (n = 241) and DART (n = 381) conditions. **d**. Same as c for cells that prefer 72 dB: 1 kHz, control: n = 207, DART: n = 323; 5 kHz, control: n = 213, DART: n = 469; 10 kHz, control: n = 367, DART: n = 344. **e**. Left, maximum responses (ΔF/F) for somatosensory stimulus responsive cells that prefer 10 PSI in control (n = 45) and DART (n = 100) conditions. Middle, same as left for cells that prefer 15 PSI: control: n = 47, DART: n = 69. Right, same as left for cells that prefer 20 PSI: control: n = 74), DART: n = 202. (auditory: 68 dB, 1 kHz preferring: 1 kHz = 5.2 ± 0.5% ΔF/F, 5 kHz = 2.0 ± 0.4% ΔF/F, 10 kHz = 1.8 ± 0.3% ΔF/F, n = 227, *P* < 0.0001; 5 kHz preferring: 1 kHz = 5.3 ± 0.6% ΔF/F, 5 kHz = 9.9 ± 0.8% ΔF/F, 10 kHz = 4.0 ± 0.4% ΔF/F, n = 211, *P* < 0.0001; 10 kHz preferring: 1 kHz = 2.4 ± 0.3% ΔF/F, 5 kHz = 2.2 ± 0.3% ΔF/F, 10 kHz = 5.4 ± 4.2% ΔF/F, n = 166, *P* < 0.0001; 72 dB, 1 kHz preferring: 1 kHz = 6.7 ± 0.5% ΔF/F, 5 kHz = 3.3 ± 0.4% ΔF/F, 10 kHz = 2.1 ± 0.2% ΔF/F, n = 184, *P* < 0.0001; 5 kHz preferring: 1 kHz = 3.4 ± 0.4% ΔF/F, 5 kHz = 6.0 ± 0.5% ΔF/F, 10 kHz = 1.6 ± 0.2% ΔF/F, n = 141, *P* < 0.0001; 10 kHz preferring: 1 kHz = 1.4 ± 0.2% ΔF/F, 5 kHz = 1.9 ± 0.2% ΔF/F, 10 kHz = 9.5 ± 0.4% ΔF/F, n = 313, *P* < 0.0001, RM ANOVA; 1 kHz, 68 dB preferring: low = 6.2 ± 0.5% ΔF/F, high = 2.7 ± 0.3% ΔF/F, n = 342, *P* < 0.0001; 72 dB preferring: low = 2.8 ± 0.2% ΔF/F, high = 5.9 ± 0.4% ΔF/F, n = 207, *P* < 0.0001; 5 kHz, 68 dB preferring: low = 8.7 ± 0.7% ΔF/F, high = 3.9 ± 0.6% ΔF/F, n = 293, *P* < 0.0001; 72 dB preferring: low = 1.8 ± 0.3% ΔF/F, high = 4.1 ± 0.3% ΔF/F, n = 213, *P* < 0.0001; 10 kHz, 68 dB preferring: low = 6.2 ± 0.4% ΔF/F, high = 1.9 ± 0.2% ΔF/F, n = 241, *P* < 0.0001; 72 dB preferring: low = 1.8 ± 0.2% ΔF/F, high = 8.6 ± 0.4% ΔF/F, n = 367, *P* < 0.0001, paired, two-tailed t-test; somatosensory: 10 PSI preferring: 10 PSI = 8.3 ± 3.1% ΔF/F, 15 PSI = 2.9 ± 1.3% ΔF/F, 20 PSI = 4.8 ± 2.8% ΔF/F, n = 45, *P* < 0.0001; 15 PSI preferring: 10 PSI = 1.3 ± 0.5% ΔF/F, 15 PSI = 4.0 ± 0.6% ΔF/F, 20 PSI = 0.7 ± 0.5% ΔF/F, n = 47, *P* < 0.0001; 20 PSI preferring: 10 PSI = 12.5 ± 3.1% ΔF/F, 15 PSI = 9.6 ± 2.3% ΔF/F, 20 PSI = 22.9 ± 4.0% ΔF/F, n = 74, *P* < 0.0001, RM one-ANOVA). (DART: auditory: 68 dB, 1 kHz preferring: 1 kHz = 16.3 ± 0.7% ΔF/F, 5 kHz = 9.4 ± 0.4% ΔF/F, 10 kHz = 13.1 ± 0.6% ΔF/F, n = 250, P < 0.0001; 5 kHz preferring: 1 kHz = 8.7 ± 0.3% ΔF/F, 5 kHz = 15.1 ± 0.5% ΔF/F, 10 kHz = 8.7 ± 0.3% ΔF/F, n = 297, P < 0.0001; 10 kHz preferring: 1 kHz = 8.0 ± 0.5% ΔF/F, 5 kHz = 7.7 ± 0.5% ΔF/F, 10 kHz = 14.8 ± 0.8% ΔF/F, n = 169, P < 0.0001; 72 dB, 1 kHz preferring: 1 kHz = 14.2 ± 0.6% ΔF/F, 5 kHz = 8.4 ± 0.5% ΔF/F, 10 kHz = 3.9 ± 0.4% ΔF/F, n = 170, P < 0.0001; 5 kHz preferring: 1 kHz = 9.3 ± 0.4% ΔF/F, 5 kHz = 18.5 ± 0.6% ΔF/F, 10 kHz = 8.7 ± 0.5% ΔF/F, n = 331, P < 0.0001; 10 kHz preferring: 1 kHz = 7.0 ± 0.3% ΔF/F, 5 kHz = 12.0 ± 0.7% ΔF/F, 10 kHz = 19.4 ± 0.7% ΔF/F, n = 224, P < 0.0001, RM ANOVA; 1 kHz, 68 dB preferring: low = 14.1 ± 0.5% ΔF/F, high = 8.1 ± 0.3% ΔF/F, n = 390, P < 0.0001; 72 Db preferring: low = 7.7 ± 0.3% ΔF/F, high = 12.1 ± 0.4% ΔF/F, n = 323, P < 0.0001; 5 kHz, 68 dB preferring: low = 15.7 ± 0.6% ΔF/F, high = 10.7 ± 0.5% ΔF/F, n = 220, P < 0.0001; 72 dB preferring: low = 9.9 ± 0.3% ΔF/F, high = 16.8 ± 0.5% ΔF/F, n = 469, P < 0.0001; 10 kHz, 68 dB preferring: low = 15.0 ± 0.45 ΔF/F, high = 5.3 ± 0.4% ΔF/F, n = 381, P < 0.0001; 72 dB preferring: low = 7.8 ± 0.3% ΔF/F, high = 17.1 ± 0.5% ΔF/F, n = 344, P < 0.0001, paired, two-tailed t-test; somatosensory: 10 PSI preferring: 10 PSI = 15.2 ± 3.1% ΔF/F, 15 PSI = 7.1 ± 2.0% ΔF/F, 20 PSI = 10.4 ± 2.9% ΔF/F, n = 100, P < 0.0001; 15 PSI preferring: 10 PSI = 4.4 ± 0.9% ΔF/F, 15 PSI = 9.9 ± 1.1% ΔF/F, 20 PSI = 5.3 ± 0.8% ΔF/F, n = 69, P < 0.0001; 20 PSI preferring: 10 PSI = 14.3 ± 1.6% ΔF/F, 15 PSI = 13.3 ± 1.5% ΔF/F, 20 PSI = 24.5 ± 2.2% ΔF/F, n = 202, P < 0.0001, RM one-ANOVA). (auditory: 68 dB, 1 kHz preferring: 5 kHz – control = 0.40 ± 0.02, DART = 0.62 ± 0.01, *P*<0.0001; 10 kHz – control = 0.38 ± 0.02, DART = 0.83 ± 0.02, *P* < 0.0001; 5 kHz preferring: 1 kHz – control = 0.48 ± 0.02, DART = 0.61 ± 0.01, *P* < 0.0001; 10 kHz – control = 0.50 ± 0.01, DART = 0.61 ± 0.01, *P* < 0.0001; 10 kHz preferring: 1 kHz – control = 0.47 ± 0.02, DART = 0.52 ± 0.02, *P* = 0.0505; 5 kHz – control = 0.42 ± 0.02, DART = 0.49 ± 0.02, *P* = 0.0149; 72 dB, 1 kHz preferring: 5 kHz – control = 0.46 ± 0.02, DART = 0.55 ± 0.02, P = 0.0028; 10 kHz – control = 0.38 ± 0.02, DART = 0.30 ± 0.02, *P* = 0.0031; 5 kHz preferring: 1 kHz – control = 0.51 ± 0.02, DART = 0.53 ± 0.01, *P* = 0.3230; 10 kHz – control = 0.41 ± 0.02, DART = 0.49 ± 0.02, *P* = 0.0120; 10 kHz preferring: 1 kHz – control = 0.23 ± 0.01, DART = 0.40 ± 0.02, P < 0.0001; 5 kHz – control = 0.29 ± 0.02, DART = 0.63 ± 0.02, *P* < 0.0001; 1 kHz, 68 dB preferring: high – control = 0.41 ± 0.01, DART = 0.60 ± 0.01, *P* < 0.0001; 72 dB preferring: low – control = 0.50 ± 0.02, DART = 0.62 ± 0.01, *P* < 0.0001; 5 kHz, 68 dB preferring: high – control = 0.48 ± 0.01, DART = 0.69 ± 0.02, *P* < 0.0001; 72 dB preferring: low – control = 0.45 ± 0.02, DART = 0.61 ± 0.01, *P* < 0.0001; 10 kHz, 68 dB preferring: high – control = 0.39 ± 0.02, DART = 0.36 ± 0.02, *P* = 0.1767; 72 dB preferring: low – control = 0.32 ± 0.01, DART = 0.50 ± 0.01, *P* < 0.0001; somatosensory: 10 PSI preferring: 15 PSI – control = 0.23 ± 0.06, DART = 0.32 ± 0.04, *P* = 0.2397; 20 PSI – control = -0.20 ± 0.28, DART = 0.45 ± 0.04, *P*= 0.0015; 15 PSI preferring: 10 PSI – control = 0.11 ± 0.14, DART = 0.37 ± 0.04, *P* = 0.0382; 20 PSI – control = 0.00 ± 0.14, DART = 0.49 ± 0.04, *P* = 0.0001; 20 PSI preferring: 10 PSI – control = 0.29 ± 0.10, DART = 0.47 ± 0.02, *P*= 0.0088; 15 PSI – control = 0.27 ± 0.10, DART = 0.47 ± 0.02, *P* = 0.0063, unpaired t-test).

**Supplemental Figure 11.**
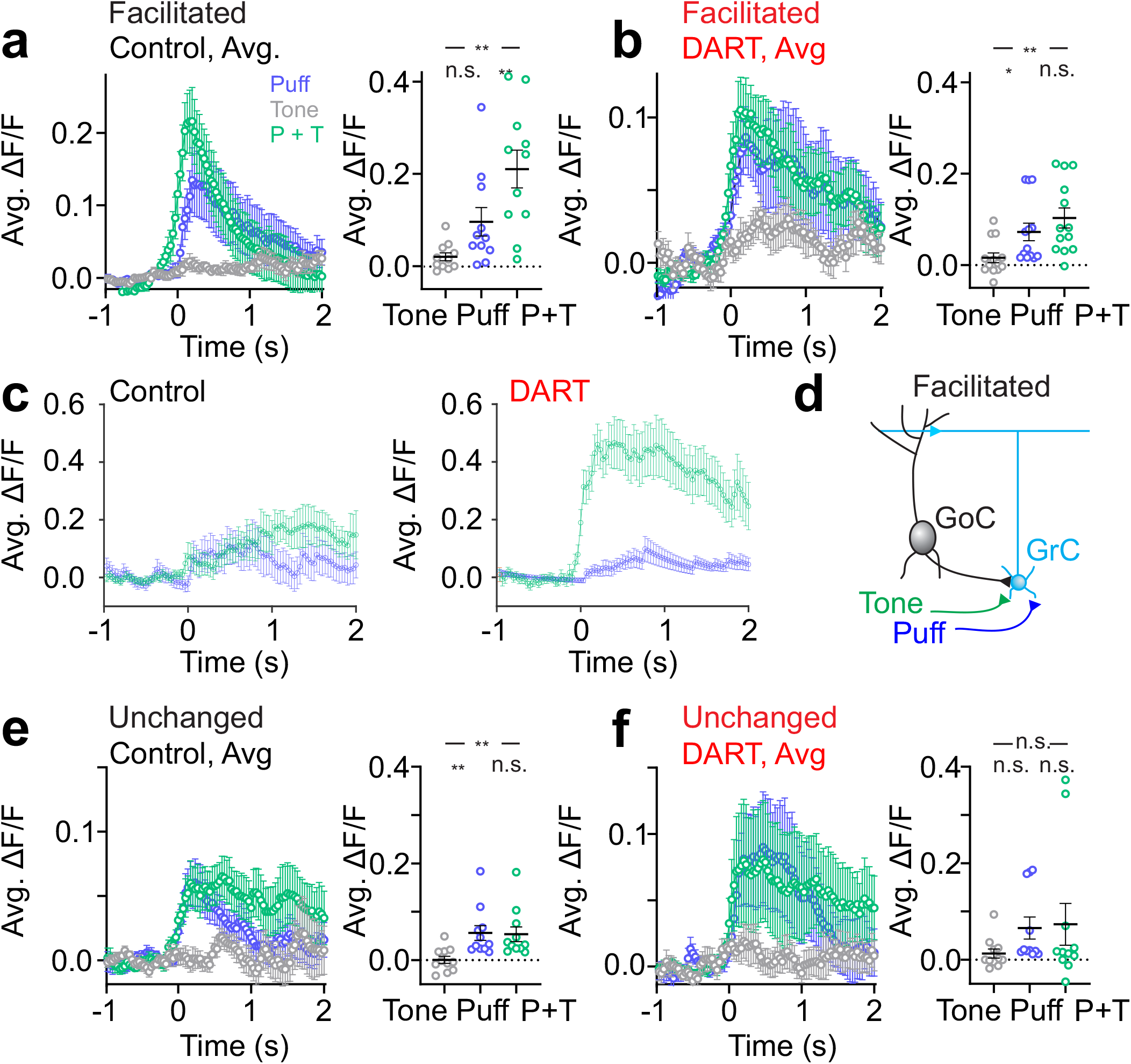
Effects of DART on cells that are facilitated or unchanged by multisensory integration. **a**. Left, average time course of responses to uni- (tone: gray; puff: blue) and multisensory (tone + puff: green) stimuli for facilitated cells (subset matched to DART, n = 13). Error is SEM across cells. Right, amplitude of responses to tone, puff and tone + puff for all facilitated cells. **b**. Same as **a**, after DART infusion. **c**. Example cell that was modestly facilitated by the tone+puff (green) as compared to the puff alone (blue) in control conditions (left), and facilitation was greatly enhanced after DART infusion (right). **d**. Proposed cerebellar circuit mediating facilitation. **e-f**. Same as **a-b**, for unchanged cells (n = 11). Note that DART does not change the integration for these cells. \**p* < 0.05, \*\**p* < 0.01.

**Supplemental Figure 12.**
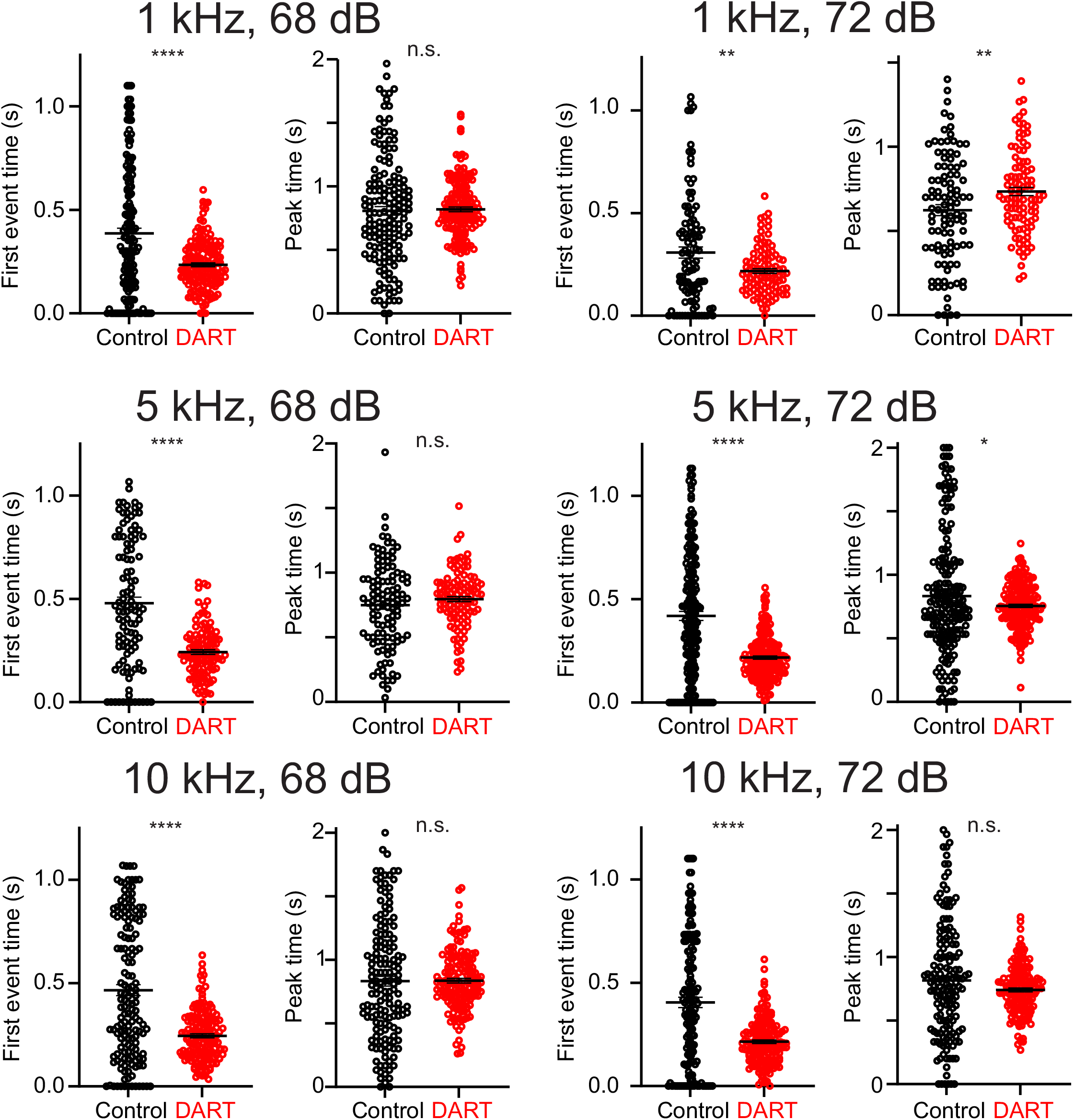
Local synaptic inhibition delays responses to all auditory stimuli tested. Mean first (left) and peak (right) event times of significant trials before (black) and after (red) DART infusion for all individual cells with significant trial responses in both conditions. Overall mean ± SEM across cells overlaid. 1 kHz, 68 dB: n = 166 cells; 1 kHz, 72 dB: n = 101 cells; 5 kHz, 68 dB: n = 110 cells; 5 kHz, 72 dB: n = 188 cells; 10 kHz, 68 dB: n = 154 cells; 10 kHz, 72 dB: n = 157 cells; 3 mice.

**Supplemental Figure 13.**
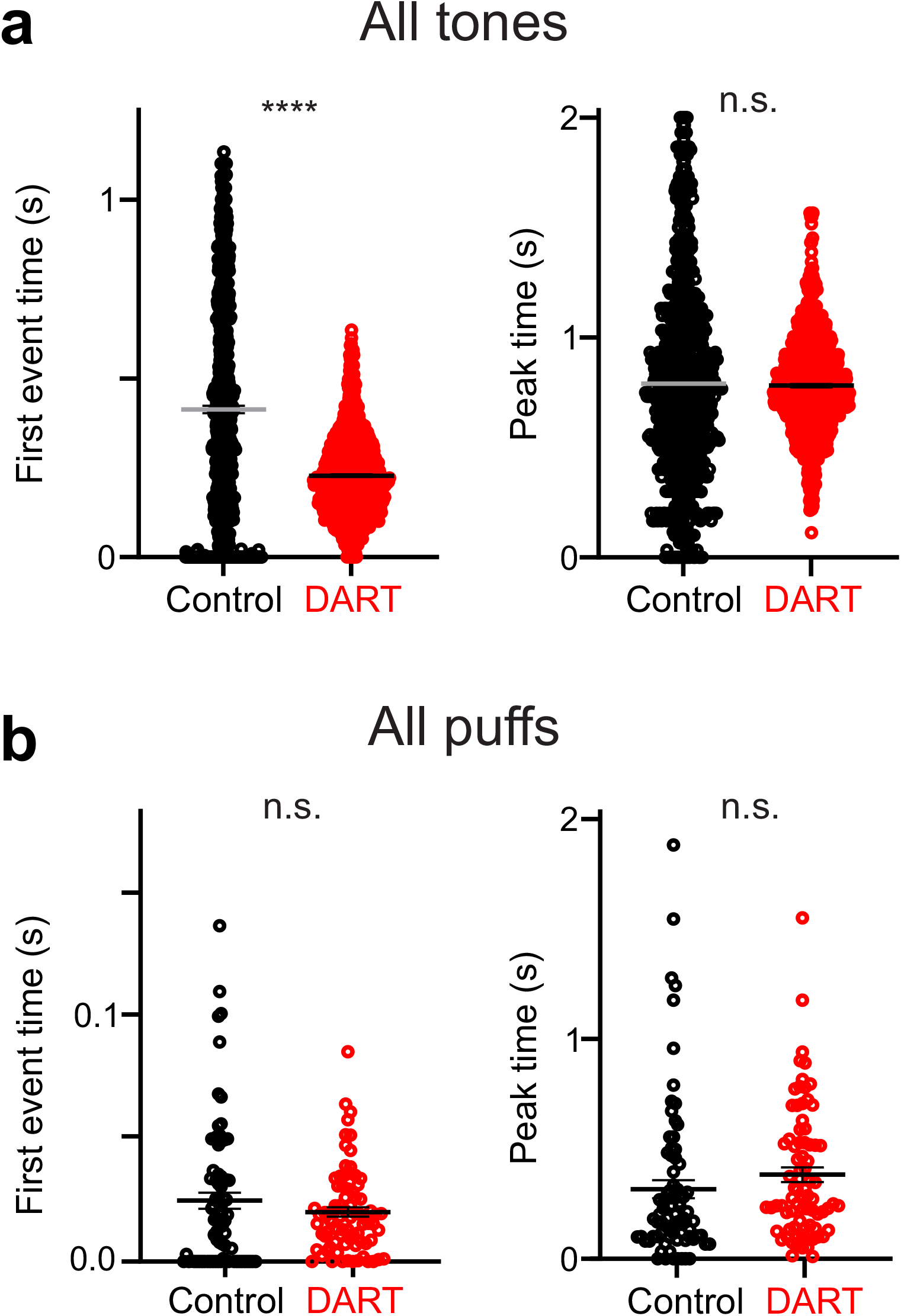
Local synaptic inhibition delays responses to auditory, but not somatosensory, stimuli. **a**. Mean first (left) and peak (right) event times of significant trials before (black) and after (red) DART infusion for all individual cells with significant trial responses to auditory stimuli in both conditions. Overall mean ± SEM across cells overlaid (n = 876). **b**. Same as **a**, for responses to air puffs (n = 104). \*\*\*\**p* < 0.0001.

**Supplemental Figure 14.**
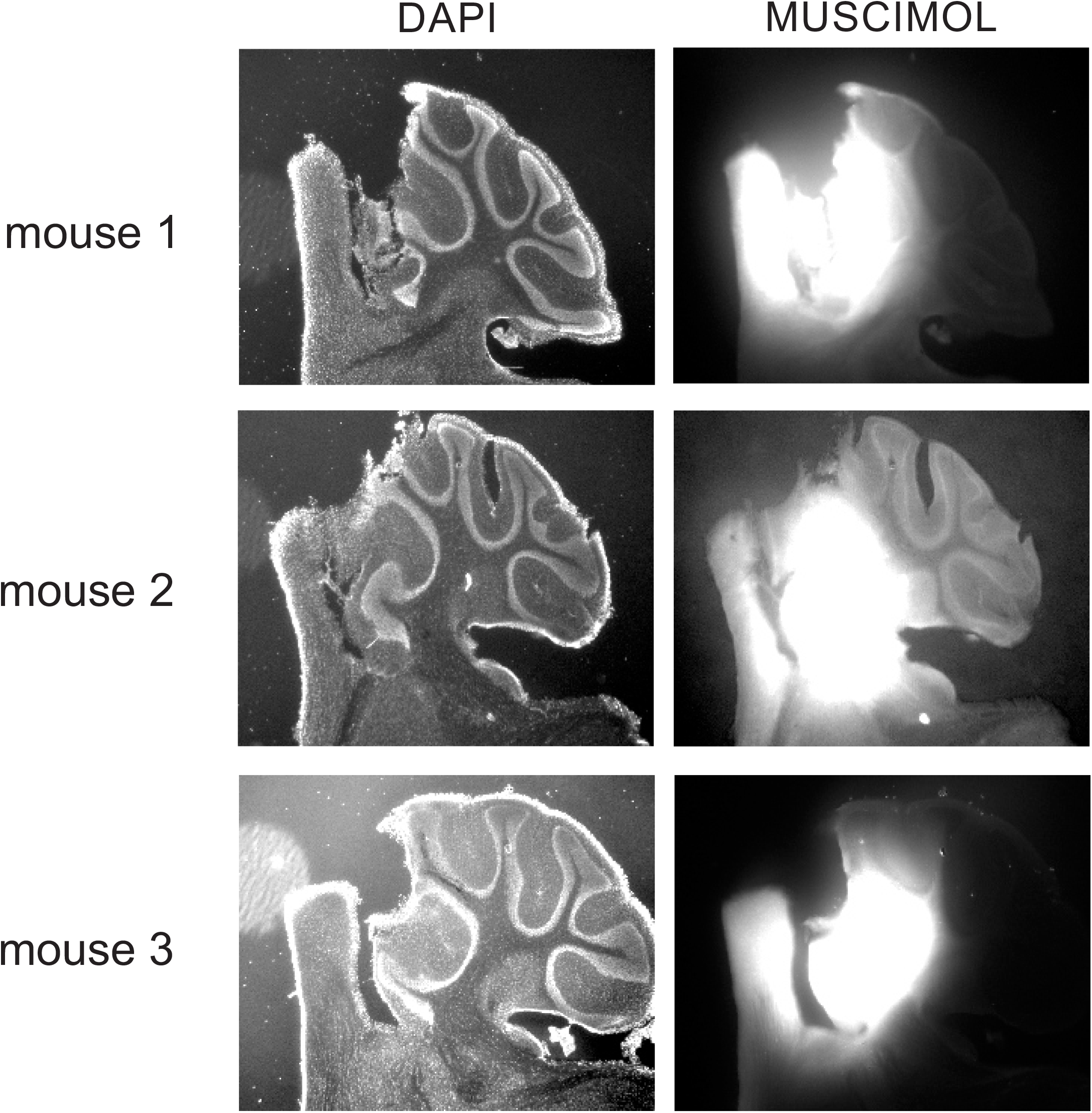
Infusion of fluorescent muscimol in conditioned mice. Example widefield images of DAPI (left) and muscimol (right) fluorescence from three mice. Muscimol injection was targeted at the floor of the primary fissure, spreading outward to include the anterior interpositus.

**Supplemental Figure 15.**
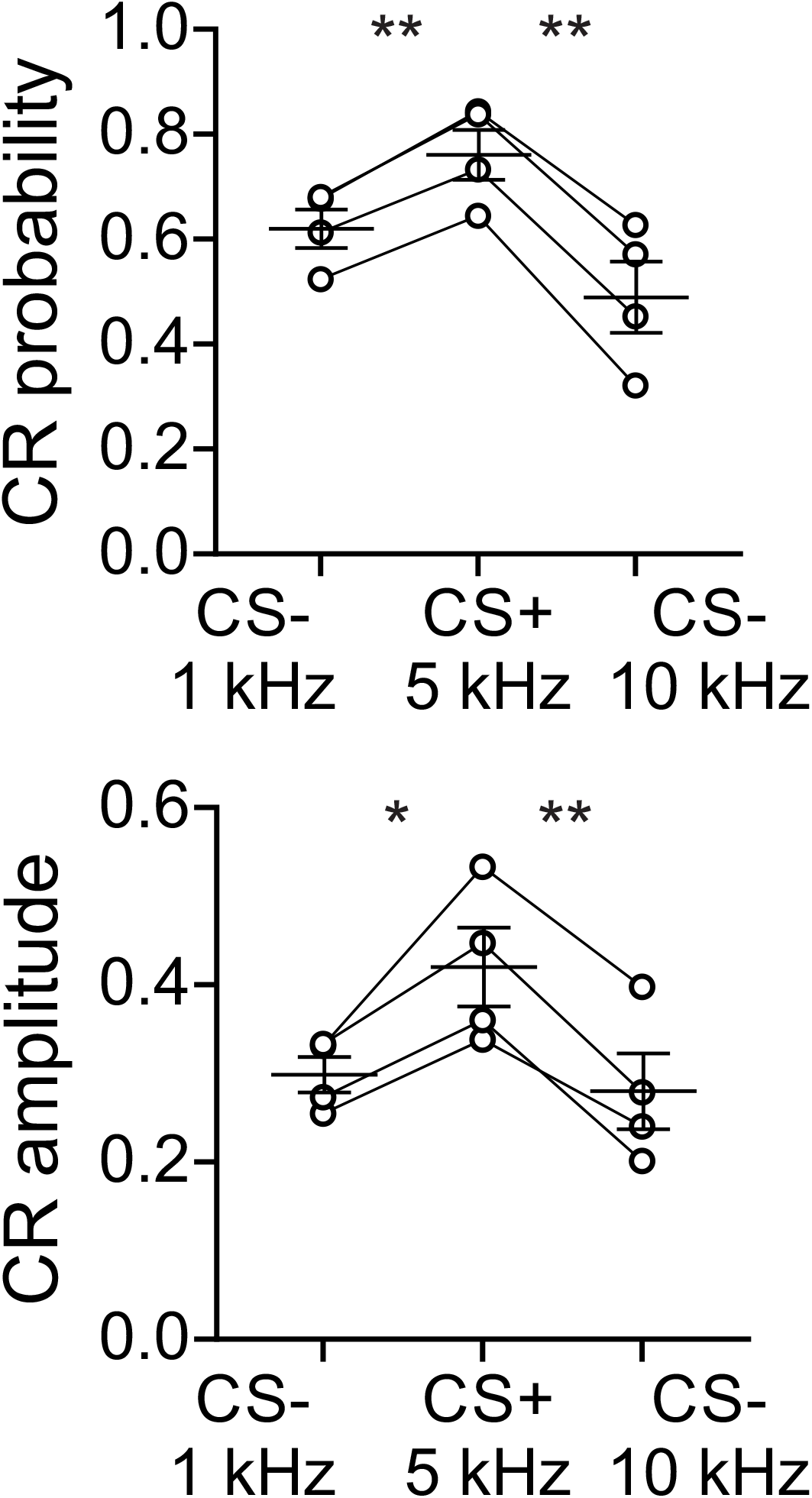
Trained mice can distinguish pure tones. Individual mean CR probability (top) and amplitude (bottom) during CS+ trials and CS- trials with either a 1 kHz or 10 kHz tone. Overall mean ± SEM across mice overlaid (n = 4 mice). \**p* < 0.05, \*\**p* < 0.01.

**Supplemental Figure 16.**
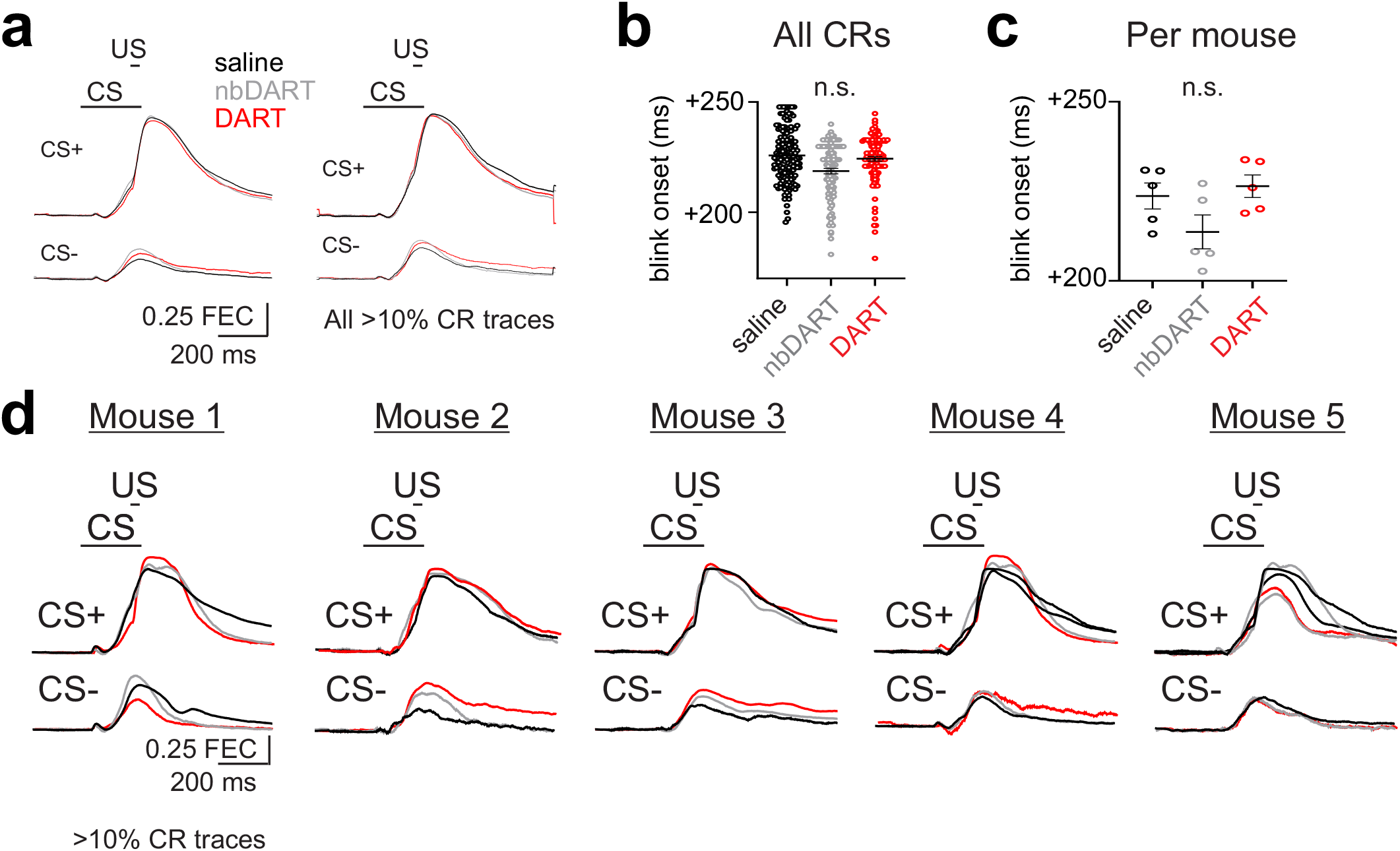
Blocking local synaptic inhibition does not change conditioned response kinematics. **a**. Average time courses of fractional eyelid closure (FEC) in response to conditioned stimuli that were either paired (CS+, top) or unpaired (CS-, bottom) with an unconditioned stimulus (US) after saline (black), nbDART (gray) or DART (red) infusion. Eyelid traces were averaged across mice **b**. Time of blink onset for all CRs in each condition for all CRs across mice (saline: n = 578, nb-DART: n = 327, DART: n = 339; RM ANOVA P < 0.0001, Tukey’s multiple comparison’s test: saline v. DART: P = 0.5837, saline v. nbDART: P < 0.0001, nbDART v. DART: P = 0.0015). Overall mean ± SEM across CRs overlaid. **c**. Summary of average time of blink onset for each mouse individually across conditions. Overall mean ± SEM across mice (n = 5) overlaid (RM ANOVA P = 0.2690). **d**. Same as **a**, for all 5 mice separately.

